# Streamlined regulation of chloroplast development in the liverwort *Marchantia polymorpha*

**DOI:** 10.1101/2023.01.23.525199

**Authors:** Nataliya E. Yelina, Eftychios Frangedakis, Zhemin Wang, Tina B. Schreier, Jenna Rever, Marta Tomaselli, Edith Forestier, Kumari Billakurthi, Sibo Ren, Yahui Bai, Julia Stewart-Wood, Jim Haseloff, Silin Zhong, Julian M. Hibberd

## Abstract

Photosynthesis in eukaryotic cells takes place in chloroplasts that develop from undifferentiated plastids in response to light. In angiosperms, after perception of light de-repression allows Elongated Hypocotyl 5 (HY5) transcription factor to initiate photomorphogenesis, and two families of transcription factors known as Golden2-like (GLK) and GATA are considered master regulators of chloroplast development. The MIR171-targeted SCARECROW-LIKE (SCL) GRAS transcription factors also impact on chlorophyll biosynthesis. The extent to which these proteins carry out conserved roles in non-seed plants is not known. Here we report in the model liverwort *Marchantia polymorpha* that GLK controls chloroplast biogenesis and HY5 shows a small conditional effect on chlorophyll content. In contrast, GATA and SCL have no detectable roles in this fundamental process. ChIP-SEQ and RNA-SEQ revealed that MpGLK regulates many photosynthetic and chloroplast development-related genes, but also has a broader set of targets than previously reported in angiosperms. This implies GLK carries out a conserved role relating to chloroplast biogenesis in land plants but also supports extensive divergence between its targets in *M. polymorpha* and flowering plants. The data support the hypothesis that regulation of chloroplast biogenesis in *M. polymorpha* is streamlined compared with angiosperms and allows us to present a core regulatory network for chloroplast biogenesis in land plants.

## Introduction

Photosynthesis sustains the majority of life on Earth and in eukaryotes is performed in organelles known as chloroplasts thought to have originated from an endosymbiotic event between a photosynthetic prokaryote and eukaryotic cell^1,2^. Chloroplasts develop in response to light from undifferentiated proplastids^3^. Key processes during chloroplast biogenesis include synthesis of chlorophyll, assembly of the thylakoid membranes and the photosynthetic apparatus and accumulation of enzymes of the Calvin Benson Bassham cycle in the chloroplast stroma^4^. Since the endosymbiotic event, the majority of genes controlling chloroplast biogenesis have transferred from plastid to nucleus and therefore proteins need to be post-translationally imported into the chloroplast^2,5^. Chloroplast biogenesis needs to be responsive to the external environment as well as the cell and so nuclear-encoded photosynthesis genes are regulated by light and hormones. Key intermediaries that allow the integration of these responses are transcription factors. Our understanding of transcription factors acting on photosynthesis and chloroplast biogenesis is based primarily on analysis of the model flowering plant *Arabidopsis thaliana* (**Figure 1A**). For example, Elongated Hypocotyl 5 (HY5), a bZIP transcription factor, acts antagonistically with Phytochrome Interacting Factors (PIFs) to activate light-regulated genes in the presence of light^6^ allowing de-etiolation and chloroplast development^7–9^. Transcription factors belonging to the GARP and GATA families also play key roles in chloroplast biogenesis^3^ and members of the SCARECROW-LIKE (SCL) GRAS family impact on chlorophyll accumulation^10^. Within the GARP superfamily, Golden2-like (GLK) are positive regulators of nuclear-encoded chloroplast and photosynthesis related genes^11–14^. In the GATA family, GATA Nitrate-inducible Carbon metabolism-involved (GNC) and Cytokinin-Responsive GATA Factor 1 (CGA1) induce genes involved in chlorophyll biosynthesis and suppress phytochrome interacting factors as well as brassinosteroid (BR) related genes to promote chloroplast biogenesis^15–18^. Moreover, GATA2 promotes photomorphogenesis by directly binding to light-responsive promoters^19^ and in the absence of light the BR-activated transcription factor BRASSINAZOLE RESISTANT1 (BZR1) represses *GATA2* expression to inhibit photomorphogenesis. Notably, a recent study showed that in *A. thaliana GLK* and *GATAs* are direct targets of HY5^20^. Lastly, three *A. thaliana* miR171 targeted SCL transcription factors redundantly regulate chlorophyll biosynthesis by repressing expression of *PROTOCHLOROPHYLLIDE OXIDOREDUCTASE C*^10^.

**Figure 1.**
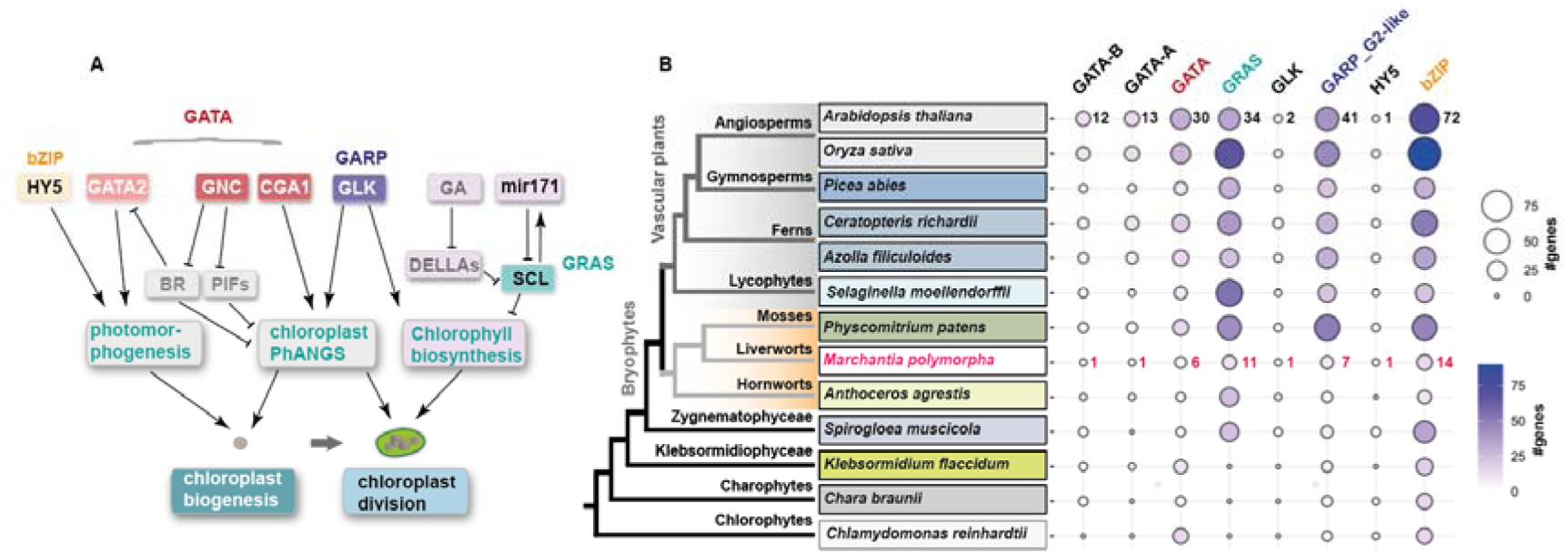
Transcription factors known to regulate chloroplast development in angiosperms. **A)** Simplified network illustrating families of transcription factors and key interacting components upstream of chloroplast biogenesis. GATA transcription factors include GATA2, GATA NITRATE-INDUCIBLE CARBON METABOLISM-INVOLVED (GNC), CYTOKININ-RESPONSIVE GATA FACTOR 1 (CGA1). GATA2 promotes photomorphogenesis in the presence of light by directly binding to light-responsive promoters. GNC and CGA1 suppress Phytochrome Interacting Factors (PIFs) and Brassinosteroid (BR) related genes, as well as promoting chloroplast biogenesis and division. In the absence of light, the BR-activated transcription factor BRASSINAZOLE RESISTANT1 (BZR1) represses *GATA2* expression. GOLDEN2-LIKE (GLK) transcription factors are positive regulators of nuclear-encoded photosynthesis related genes. SCARECROW-LIKELJ(SCL) GRAS transcription factors are negatively regulated by miR171 and GA-DELLA signalling to control chlorophyll biosynthesis. Elongated Hypocotyl 5 (HY5), a bZIP transcription factor that plays a primary role in de-etiolation and chloroplast development. **B)** Phylogenetic relationships for major lineages of land plants and green algae (Li et al. 2020). Gene numbers shown for transcription factor families and subfamilies in (A).

Land plants evolved from aquatic green algae and approximately 500 million years ago (MYA) diverged into two major monophyletic clades comprising vascular plants (angiosperms, gymnosperms, ferns and lycophytes) and bryophytes (hornworts, liverworts and mosses) (**Figure 1B**)^21–23^. Bryophytes are of particular importance to infer more accurately the ancestral state of a trait. Whilst earlier evolutionary hypotheses proposed they represent a collection of paraphyletic lineages^24^ it is now thought that liverworts and mosses form a monophyletic group that split from hornworts approximately 400-500 MYA^21,23,25^. This revised phylogeny supports the notion that traits found in the common ancestor of land plants could have diversified not only within the vascular plant clade but also in the three deeply divergent bryophytes groups. Thus, studying representative species from more than one bryophyte clade provides insight into the likely ancestral state. Despite our detailed knowledge of chloroplast biogenesis in angiosperms, with the exception of GLK function in the moss *Physcomitrium patens*^26^ our understanding of how chloroplast biogenesis is controlled and evolved remains unclear. *M. polymorpha* has a small and well-annotated genome, key steps in its development are easily accessible for observation, and an extensive set of genetic manipulation tools are available^27,28^. We therefore selected the model liverwort *Marchantia polymorpha* to investigate processes underpinning the evolution of chloroplast biogenesis.

We first performed phylogenetic analysis to identify the *M. polymorpha* homologs of HY5, GLK, GATAs and the SCL-MIR171, and generated knock-out mutants to test whether each component impacts on chloroplast biogenesis in *M. polymorpha*. We found that only HY5 and GLK are important for this process in *M. polymorpha,* but Mp*GATA* and Mp*SCL* are not. When *GLK* was over-expressed in *M. polymorpha* the abundance of transcripts derived from genes associated with chlorophyll biogenesis and Photosystems I and II were impacted and ChIP-SEQ confirmed that GLK can bind promoters of those genes. Intriguingly, many MpGLK targets are distinct from those documented in flowering plants. We conclude GLK function is conserved between *M. polymorpha* and angiosperms but that other regulators such as *GNC/CGA1, GATA2* and *SCL* have no detectable impact on chloroplast development. Thus, in comparison to angiosperms, the pathway controlling chloroplast biogenesis in *M. polymorpha* is more streamlined.

## Results

### *Identification of M. polymorpha* homologs of HY5, GLK, *GATA* and *SCL* from angiosperms

While a HY5 homolog has previously been identified in *M. polymorpha*^29^, there is currently no systematic analysis of GATA, GLK and SCL in this context. We therefore used phylogenetic analysis to search for *GATA* (*GNC*, *CGA1* and *GATA2*), *SCL* and *GLK* orthologs in *M. polymorpha*. To do so we examined twenty-one representative species from the seven main groups of land plants as well as four green algae for which high quality genome assemblies are available (**Figure 1B, Figure S1A**).

*GNC*, *CGA1* and *GATA2* belong to the GATA superfamily of transcription factors that comprise a family of zinc finger proteins present in all eukaryotes^30,31^ (**Figure 2A and Figure S1B&C**). *M. polymorpha* has a total of six *GATA* genes, among which we identified *Mp7g03490* (annotated as Mp*GATA4*) and *Mp1g03950* (annotated as Mp*GATA2*) as single orthologs of *GNC/CGA1* and *GATA2* respectively (**Figure 1B, Figure S1B, S1C, S2 S3A, and Dataset S1**). *SCL* is a member of the GRAS family of transcription factors which have a conserved C-terminal domain (**Figure 1B and Figure 2A**). *A. thaliana AtSCL6, AtSCL22* and *AtSCL27* redundantly control chlorophyll biosynthesis and are regulated by MIR171. The *M. polymorpha* genome encodes eleven GRAS transcription factors. *Mp8g03980* has been annotated as Mp*GRAS10* and is located in a sister clade to *A. thaliana* At*SCL6*, At*SCL22* and At*SCL27*^32^. Based on conservation of its miR171-mediated regulation we refer to *Mp8g03980* as Mp*SCL*. Lastly, *GLK* genes belong to the GARP (GOLDEN2-LIKE, ARR-B, Psr1) family of transcription factors^33^ (**Figure 1B, 2A, Figure S1D and S3B**)^13^. Our phylogenetic analysis identified *Mp7g*09740 (annotated as Mp*GARP8*) as a single ortholog of *GLK* due to presence of the characteristic C-terminal GOLDEN2 C-terminal (GCT) box^13^ (**Figure S1D,E and Dataset S1**). In summary, our analysis indicates that *M. polymorpha* contains single orthologs of *GNC/CGA1*, *GATA2*, *SCL* and *GLK* that we hereafter refer to as Mp*GATA4*, Mp*GATA2*, Mp*SCL* and Mp*GLK* respectively.

**Figure 2.**
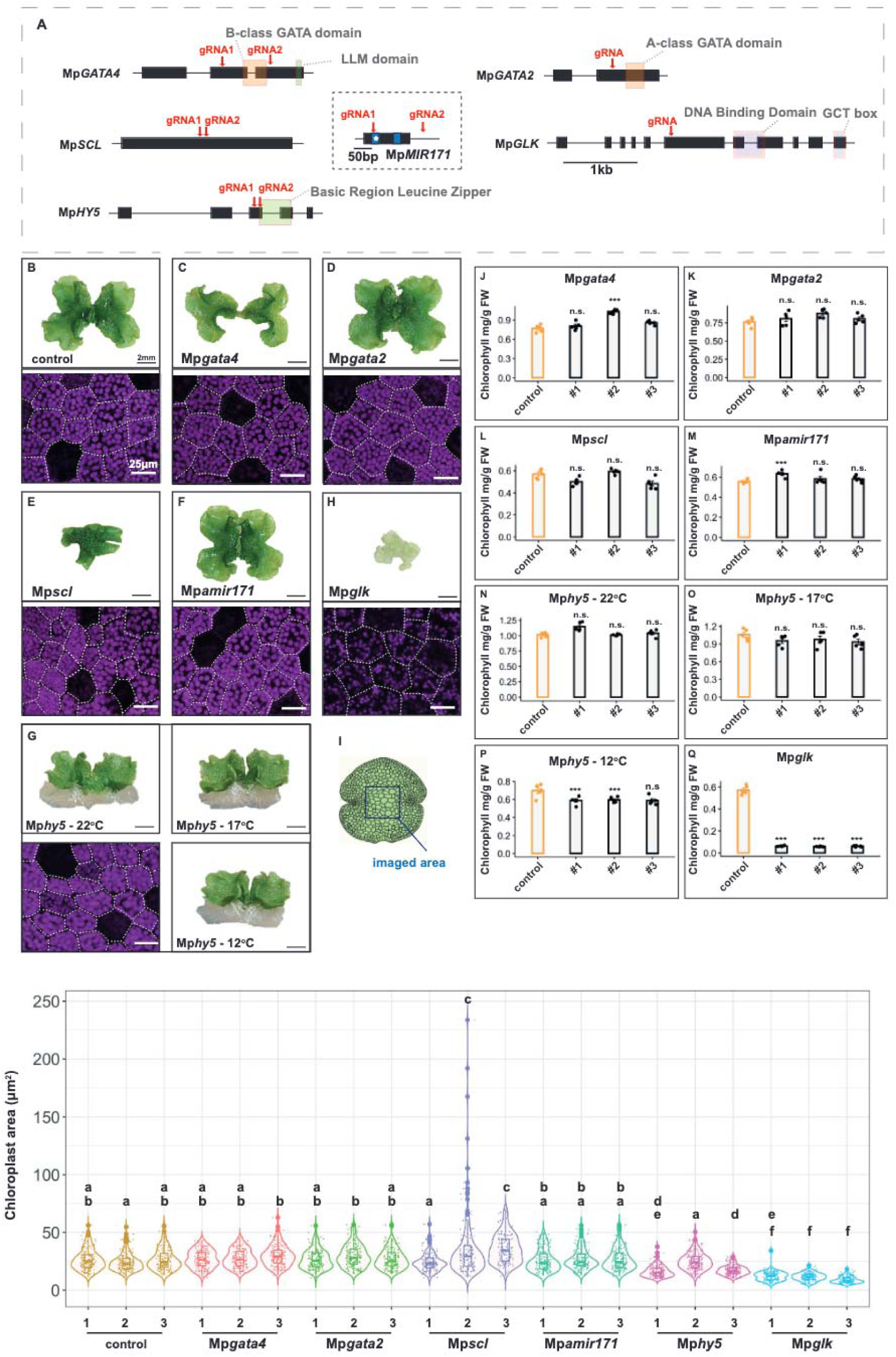
Knockout mutants for *HY5, GLK, GATA* and *SCL*. **A)** Schematic representation of Mp*GLK,* Mp*GATA4,* Mp*GATA2,* Mp*MIR171,* Mp*SCL* and Mp*HY5* gene structure showing exons as black rectangles. Canonical domains for each family are shown as coloured boxes. Genomic locus of Mp*MIR171* represented as a black rectangle, miR171* and miR171 shown as blue rectangles, miR171* indicated with a white star. Guide RNA positions for CRISPR/Cas9 gene editing shown as red arrows. **B-G)** Top: Representative images of wild type, Mp*gata4*, Mp*gata2*, Mp*scl*, Mp*mir171,* Mp*hy5* and Mp*glk* mutants. Bottom: Representative confocal microscopy images of wild type, Mp*gata4*, Mp*gata2*, Mp*scl*, Mp*mir171* and Mp*glk* mutants. Chlorophyll autofluorescence is shown in magenta and cell boundaries marked with dashed lines. Scale bars: 1 cm. **I)** Schematic of a gemmae with dark blue square indicating imaged area, **J-Q)** Barplots of chlorophyll content for the Mp*gata4*, Mp*gata2*, Mp*scl*, Mp*mir17*1, Mp*hy5* and Mp*glk* mutant plants. Individual values are shown with dots. Error bars represent the standard deviation of the mean *n* = 5. Asterisks indicate statistically significant difference using a two-tailed *t*-test *P* ≤ 0.0001 (***). **R)** Boxplot showing chloroplast size range in control versus Mp*gata4*, Mp*gata2*, Mp*scl*, Mp*mir171,* Mp*hy5* and Mp*glk* mutants. Box and whiskers represent the 25 to 75 percentile and minimum-maximum distributions of the data. Letters show statistical ranking using a *post hoc* Tukey test (with different letters indicating statistically significant differences at P<0.01). Values indicated by the same letter are not statistically different. n = 150.

### Knock-out mutant analysis of *HY5, GLK, GATA* and *SCL*

To test whether Mp*GATA4*, Mp*GATA2*, the Mp*SCL-MIR171* module, Mp*HY5* and Mp*GLK* orthologs control chloroplast biogenesis in *M. polymorpha* we used CRISPR/Cas9 editing to generate knock-out mutant alleles for each (**Figure 2A**). Plants transformed with the same CRISPR/Cas9 containing vector but without a guide RNA (gRNA) sequence were used as ‘no guide RNA’ controls. Each mutant line was clonally propagated through one gemmae generation to obtain isogenic plants, and for each targeted gene three independent lines were selected for analysis. In all protein coding genes, mutations led to premature stop codons (**Figure S4**) and Mp*mir171* mutants had a 139 bp deletion and 1 bp insertion resulting in a deletion of the entire Mp*MIR171* gene (**Figure S4D**).

In Mp*gata4* mutants, gametophyte morphology was perturbed with narrower thallus lobes (**Figure 2B, C**). Compared with controls, chlorophyll content was ∼10% higher in one of the three lines examined (**Figure 2J**), a response opposite to the expected reduction if *GATA4* plays a conserved role. We also analysed chloroplast morphology via confocal laser scanning microscopy (**Figure 2I**). In Mp*gata4* mutants, chloroplast size in cells of the central part of gemmae (excluding the rhizoid precursors) was similar to that of controls (**Figure 2C, R**). Mp*gata2* mutants did not show any morphological or developmental phenotypes and had similar chlorophyll levels compared with ‘no guide RNA’ controls (**Figure 2B, D, K**). Chloroplast size in Mp*gata2* mutants were also not perturbed (**Figure 2D, R**). Mp*scl* mutants showed altered morphology with thalli being ‘stunted’ with inward curling edges (**Figure 2B, E**). Despite chlorophyll levels being not statistically different from controls (**Figure 2L**) in two of the three Mp*scl* mutant lines we observed an increase in the size of the chloroplasts, which was statistically significant in some cases (**Figure 2E, R**). In contrast, Mp*mir171* mutants were indistinguishable from controls (**Figure 2B, F, M, R**). Mp*hy5* gametophyte morphology was also perturbed with thallus lobes being more erect (**Figure 2G**). Compared with controls, at 22 °C, chloroplast size was ∼15% smaller in two of the three Mp*hy5* lines (**Figure 2G, R**). In *A. thaliana hy5* mutants show a conditional phenotype with greater impact at lower temperature^34^. To test whether HY5 also has a conditional phenotype in *M. polymorpha* we grew plants at 22 °C, 17 °C or 12 °C. At 22 °C and 17 °C mutants and controls had similar chlorophyll content (**Figure 2G, N, O**), but at 12 °C a small but statistically significant reduction in chlorophyll was evident (**Figure 2G, P**). Several sesquiterpenes were more abundant in Mp*hy5* compared with controls (**Dataset S2**). We conclude that *hy5* mutants from *M. polymorpha* and *A. thaliana* have similar temperature dependent phenotype relating to chloroplast biogenesis, but in *M. polymorpha* the protein also appears to impact on terpenoid accumulation, a role distinct from that previously reported for flavonoid biosynthesis^29^.

Mp*glk* mutants had an obvious pale green phenotype (**Figure 2B, H**). Mutations that either caused a deletion of the GARP DNA-binding domain or introduced a premature stop codon immediately upstream of it, resulted in a more severe phenotype (we refer to these as ‘strong’ alleles) than mutations in the 5′ end of the Mp*GLK* gene (‘weak’ alleles) (**Figure S5A-F**). In the ‘strong’ Mp*glk* mutant alleles chlorophyll content was reduced by ∼90% compared with controls and chloroplasts were smaller (**Figure 2H, Q, R**). Mp*glk* mutants also showed morphological changes with narrower thallus lobes and upward curling lobe edges (**Figure 2H**). However, they were still able to grow and produce gemmae and reproductive organs (**Figure S5G**) without supplemental carbon. Consistent with confocal laser scanning microscopy, electron microscopy confirmed Mp*glk* mutants had smaller chloroplasts compared with controls and ultrastructure was perturbed (**Figure 3A, B**). Specifically, mutant chloroplasts had fewer thylakoid membranes with reduced granal stacking (**Figure 3A, B and Figure S5H**). Treatment with di-chlorophenyl di-methyl urea (DCMU) which inhibits the photosynthetic electron transport chain^35^ caused a substantial reduction in the chlorophyll fluorescence parameter F_v_/F_m_ in mutants indicating that their photosynthetic apparatus was functional (**Figure 3C, D**). This is consistent with the ability of ‘strong’ Mp*glk* alleles to grow under standard conditions without a carbon supplement.

**Figure 3.**
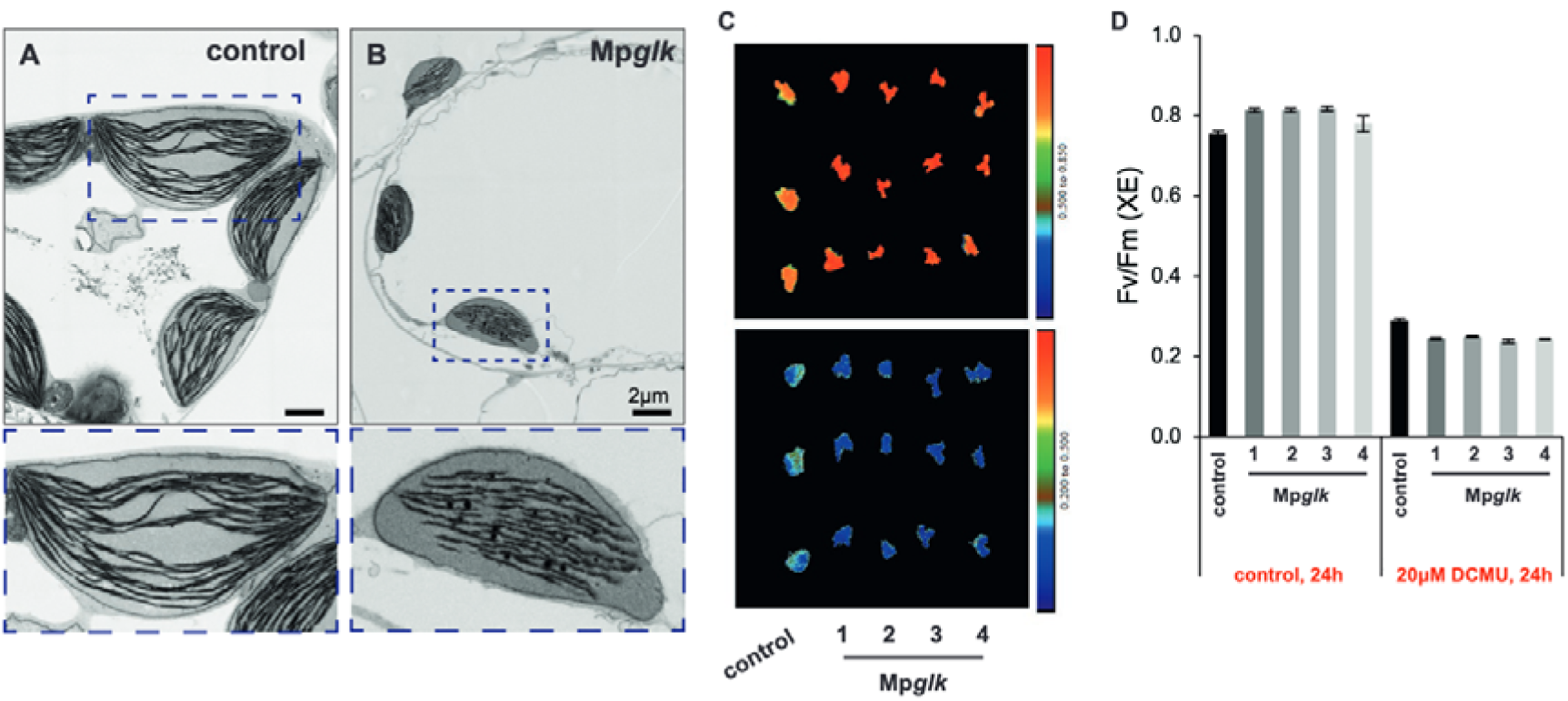
Mp*glk* controls chloroplast size and ultrastructure. **A-B)** Scanning electron micrographs of chloroplasts from control (A) and Mp*glk* mutants (B). Dashed boxes highlight single chloroplasts that are shown in the insets at higher magnification (bottom panels). Scale bars: 2 μm. **C)** Chlorophyll fluorescence images of maximum quantum efficiency of Photosystem II (*Fv/Fm*) of untreated (top) and 24 h DCMU-treated (bottom) control and Mp*glk* mutants. **D)** *Fv/Fm* measured in DCMU-treated and untreated control and Mp*glk* mutants. Bars represent mean ± standard error from *n* = 3 plants per genotype.

In summary, the absence of functional Mp*GATA4* and Mp*GATA2* genes did not appear to affect chlorophyll biosynthesis or chloroplast biogenesis in *M. polymorpha*. As *GNC/CGA1* mutants of *A. thaliana* contain reduced chlorophyll^36^ our data argue against functional conservation between *A. thaliana* GNC/CGA1 and Mp*GATA4* in regulating chloroplast biogenesis. Similarly, we found a lack of conservation between the function of At*GATA2* and Mp*GATA2*. Moreover, despite conservation of a MIR171 target site in the Mp*SCL* mRNA, MpSCL does not appear to control chlorophyll content. Again, this is in contrast with *A. thaliana* where triple At*scl6,scl22,scl27* mutants showed increased chlorophyll, and At*MIR171* mimic lines, which are functional equivalents of At*mir171* knock-outs, had reduced chlorophyll^10,37^. Although chloroplast sizes were unaltered in Mpg*ata2*, Mp*gata4* or Mp*mir171,* we found that two of the three Mp*scl* lines assessed had larger chloroplasts. In contrast, as in *A. thaliana*, Mp*hy5* mutants showed a conditional effect with lower chlorophyll at low temperature, and Mp*glk* mutants accumulated less chlorophyll and had smaller chloroplasts than wild type. Collectively our results imply a strongly conserved function for GLK in chloroplast biogenesis in *M. polymorpha* and suggest that the conditional role of HY5 may be also conserved.

### Limited interplay between GATA4, GATA2, SCL, HY5 and GLK in *M. polymorpha*

It is possible that redundancy or compensatory responses mask effects on chloroplast biogenesis in loss of function alleles for MpGATA4, MpGATA2, MpSCL and MpHY5. To test this, we generated Mp*glk,gata4*, Mp*glk,gata2,* Mp*glk,scl* and Mp*glk,hy5* double mutants (**Figure 4A-E and Figure S5I**). None were noticeably paler than the single Mp*glk* mutant (**Figure 4A-E**) and chlorophyll content was similar (**Figure 4F**). In Mp*glk,gata4,* Mp*glk,scl* and Mp*glk,hy5* double mutants, thallus morphology was perturbed in a similar manner to that seen in single Mp*gata4,* Mp*scl* and Mp*hy5* mutants. Chloroplast size in double Mp*glk,gata4* mutants was also comparable to single Mp*glk* mutants (**Figure 4G**). Moreover, over-expression of MpGATA4 in Mp*glk* did not rescue the pale Mp*glk* phenotype (**Figure 4H and S5J**). In *A. thaliana*, HY5 regulates *GLK* and *GNC/CGA1* ^38^ and so we asked if this was also the case for the homologs in *M. polymorpha*. We therefore subjected 6-day-old Mp*hy5* and control gemmalings to 48 hours of darkness, followed by exposure to 1.5, 3 and 6 hours of light. However, Mp*GLK* and Mp*GATA4* expression was comparable in Mp*hy5* and controls, providing no evidence for Mp*GLK* and Mp*GATA4* being targeted by MpHY5 (**Figure 4I**). In summary, the data provide no evidence for functional redundancy between Mp*GATA4*, Mp*GATA2,* Mp*SCL*, Mp*HY5* and Mp*GLK*.

**Figure 4.**
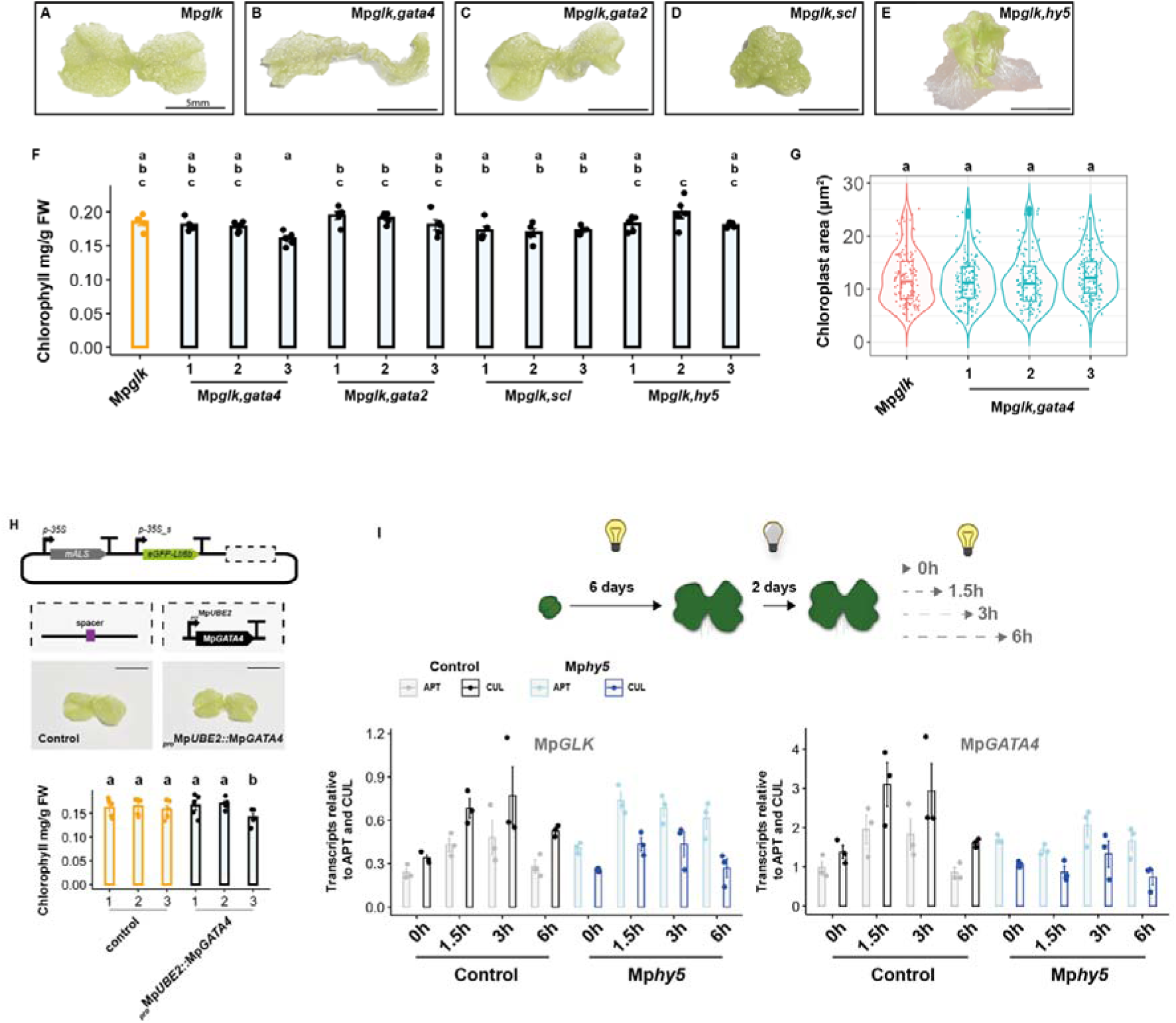
*GATA4, GATA2, SCL* and *HY5* are not epistatic with *GLK* for chloroplast biogenesis in *M. polymorpha*. **A-E)** Representative images of Mp*glk,* Mp*glk,gata4,* Mp*glk,gata2,* Mp*glk,scl* and Mp*glk,hy5* double mutants. **F)** Chlorophyll content in Mp*glk,* Mp*glk,gata4,* Mp*glk,gata2,* Mp*glk,scl* and Mp*glk,hy5* double mutants. Letters show statistical ranking using a *post hoc* Tukey test, different letters indicating statistically significant differences at P<0.01. Values indicated by the same letter are not statistically different, n=5. **G)** Chloroplast area in Mp*glk,* and Mp*glk,gata4.* Letters show statistical ranking using a *post hoc* Tukey test, different letters indicating statistically significant differences at P<0.01. Values indicated by the same letter are not statistically different, n=150. **H)** Top: Schematic representation of Mp*GATA4* overexpression construct. Bottom: Representative images of Mp*glk* mutants and Mp*glk* mutant overexpressing Mp*GATA4*. Scale bar represents 5 mm. Bottom: Chlorophyll content in Mp*glk* mutant and Mp*glk* mutant overexpressing Mp*GATA4*. Letters show statistical ranking using a *post hoc* Tukey test, different letters indicating statistically significant differences at P<0.01. Values indicated by the same letter are not statistically different, n=5. **I)** Top: Schematic of experimental design to test the effect of lack of functional MpHY5 on Mp*GLK* and Mp*GATA4* expression in response to light. Bottom: quantitative polymerase chain reaction (qPCR) for analysis of Mp*GLK* and Mp*GATA4* expression in Mp*hy5* mutants in response to light. *ADENINE PHOSPHORIBOSYL TRANSFERASE 3* (*APT*) and *CULLIN 1* (*CUL*) were used as housekeeping gene controls (Saint-Marcoux et al., 2015).

### Mp*GLK* is sufficient to activate chloroplast biogenesis

To determine whether homologs of Mp*GATA4,* Mp*GATA2,* Mp*SCL* and Mp*GLK* are sufficient to activate chloroplast biogenesis we generated over-expression lines driven by the strong constitutive ubiquitin-conjugating enzyme E2 gene promoter (Mp*UBE2*). To facilitate analysis GFP was used to mark the plasma membrane (**Figure 5A**). No differences in chlorophyll or thallus morphology between plants with and without the plasma membrane marker were evident (**Figure S5K**). qPCR confirmed that each transgene was over-expressed (**Figure S5L**). No differences in chlorophyll content or chloroplast size were detected between controls and over-expressing lines of Mp*GATA4,* Mp*GATA2* or Mp*SCL* (**Figure 5B-F, I, J**). Since in *A. thaliana, GATA2* over-expression led to constitutive photomorphogenesis ^19^ we placed gemmae over-expressing Mp*GATA2* in the dark alongside controls and examined if there were any differences in growth after a week. Over-expressors and controls were indistinguishable (gemmae did not germinate) (**Figure S6A**). However, plants in which Mp*GLK* was over-expressed were darker green and displayed stunted growth (**Figure 5G**). In fact, lines over-expressing Mp*GLK* contained up to ∼4 times higher chlorophyll content, and chloroplast size was significantly increased (**Figure 5I, J**). Interestingly, we observed an increase in chloroplast size in normally non-photosynthetic rhizoid precursor and oil body cells compared with controls (**Figure 5H**). Moreover, the 3’UTR of Mp*GLK* reduced greening that was potentiated by its over-expression (**Figure S6B-H**). We conclude that only Mp*GLK* is sufficient to activate chloroplast biogenesis.

**Figure 5.**
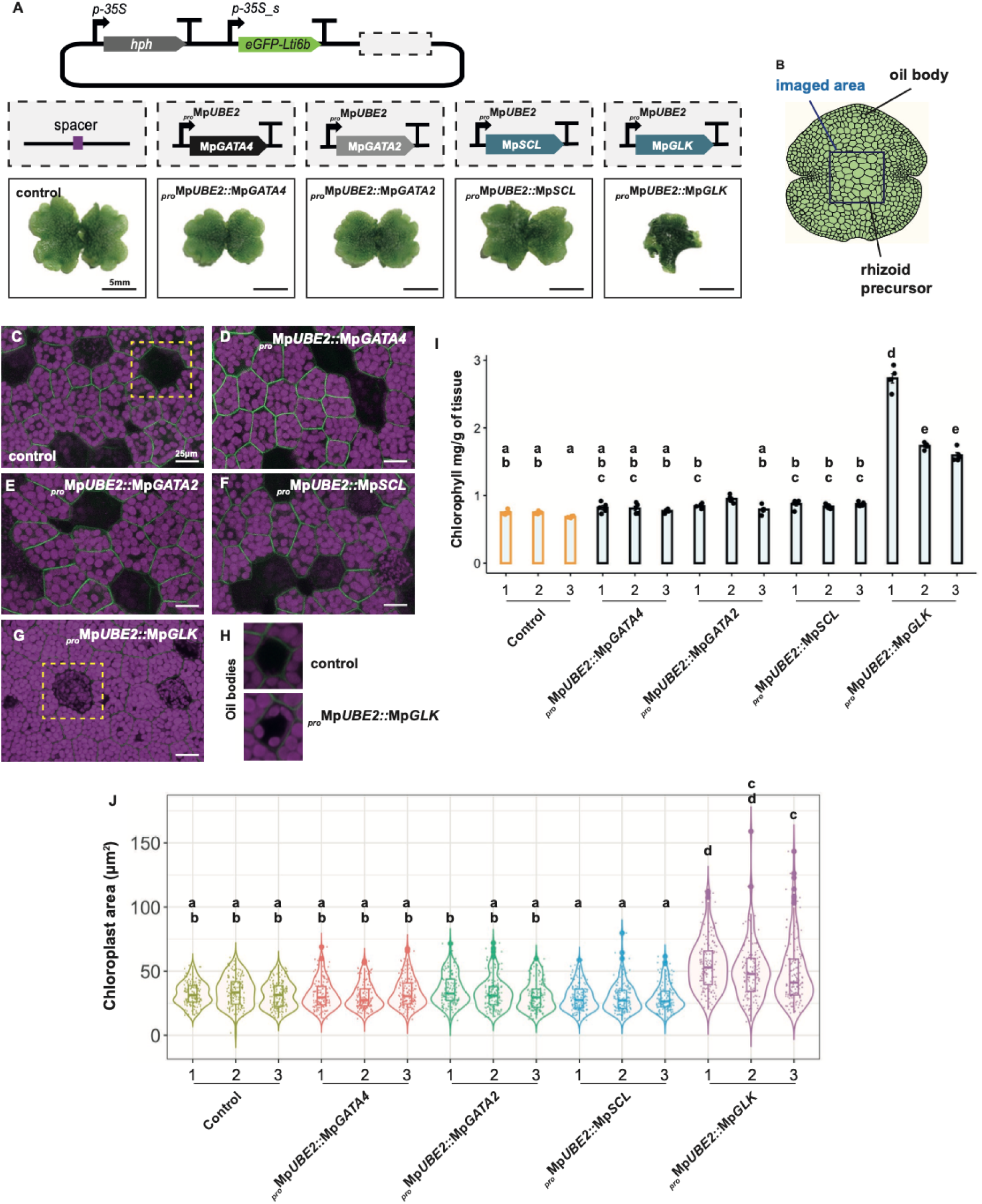
Mp*GLK* overexpression increases chlorophyll content and chloroplast size. **A)** Top: Schematic representation of constructs used to overexpress Mp*GATA4,* Mp*GATA2,* Mp*SCL* and Mp*GLK.* Bottom: Representative images of controls and plants overexpressing Mp*GATA4,* Mp*GATA2,* Mp*SCL* and Mp*GLK*. Scale bar represents 5 mm. **B)** Schematic of a gemmae; a dark blue square indicates imaged area, black lines indicate approximate positions of rhizoid precursor and oil body cells. **C-G)** Representative confocal microscopy images of gemmae from controls and plants overexpressing Mp*GATA4,* Mp*GATA2,* Mp*SCL* and Mp*GLK*. Chlorophyll autofluorescence shown in magenta and plasma membrane marked with eGFP in green. Rhizoids precursors highlighted with dashed squares. Scale bars, represent 25 μm. **H)** Representative confocal microscopy images of oil body cells in control and Mp*GLK-CDS* overexpressing lines. **I)** Chlorophyll content in controls and overexpressing lines for Mp*GATA4,* Mp*GATA2,* Mp*SCL* and Mp*GLK.* Letters show statistical ranking using a *post hoc* Tukey test (different letters indicating statistically significant differences at P<0.01). Values indicated by the same letter are not statistically different, n=5. **J)** Chloroplast area in controls and lines overexpressing Mp*GATA4,* Mp*GATA2,* Mp*SCL* and Mp*GLK.* Letters show statistical ranking using a *post hoc* Tukey test (different letters indicating statistically significant differences at P<0.01). Values indicated by the same letter are not statistically different, n=150.

### GLK in *M. polymorpha* regulates thylakoid associated photosynthetic components and chlorophyll biosynthesis

Although there were limited effects of knocking out *HY5*, *GATA4* and *SCL* on chloroplast biogenesis in *M. polymorpha* we undertook RNA sequencing (RNA-SEQ) on these lines along with those we generated for Mp*GLK*. Knocking out Mp*SCL*, Mp*GATA4* and Mp*HY5* had a moderate effect with 243, 332 and 160 genes being downregulated respectively (*padj*-value =<0.01, LFC ≥ 1-fold) (**Figure S7A-C, Dataset S3**). In loss of function mutants for Mp*GLK,* 1065 genes showed a reduction in transcript abundance compared with controls (*padj*-value ≤0.01, LFC ≥ 1-fold) (**Figure S6D, Dataset S3**). Changes to transcript abundance that were common between genotypes were limited with the largest overlap (85 genes) being detected for Mp*glk* and Mp*gata4* mutants (**Figure 6A**). In all three genotypes Gene Ontology^39^ analysis showed that oxidative stress as well as hydrogen peroxide catabolic processes were most impacted (**Figure 6B**). However, in Mp*glk* photosynthesis and chlorophyll biosynthesis process were also affected (**Figure 6B**) and analysis of the effects of loss of Mp*SCL,* Mp*GATA4,* Mp*HY5* and Mp*GLK* function on chlorophyll biosynthesis and canonical photosynthesis genes confirmed the greatest effect in Mp*glk*. For example, in Mp*scl* only one chlorophyll biosynthesis gene, *CAO*, was upregulated (Log2FC∼0.58). In Mp*hy5* and Mp*gata4* mutants, there was a moderate reduction in transcript abundance (Log2FC∼-0.3-0.7) of thirteen and five chlorophyll biosynthesis genes respectively (**Figure 6C**). In contrast, Mp*GLK* mis-expression affected seventeen of the nineteen chlorophyll biosynthesis genes.

**Figure 6.**
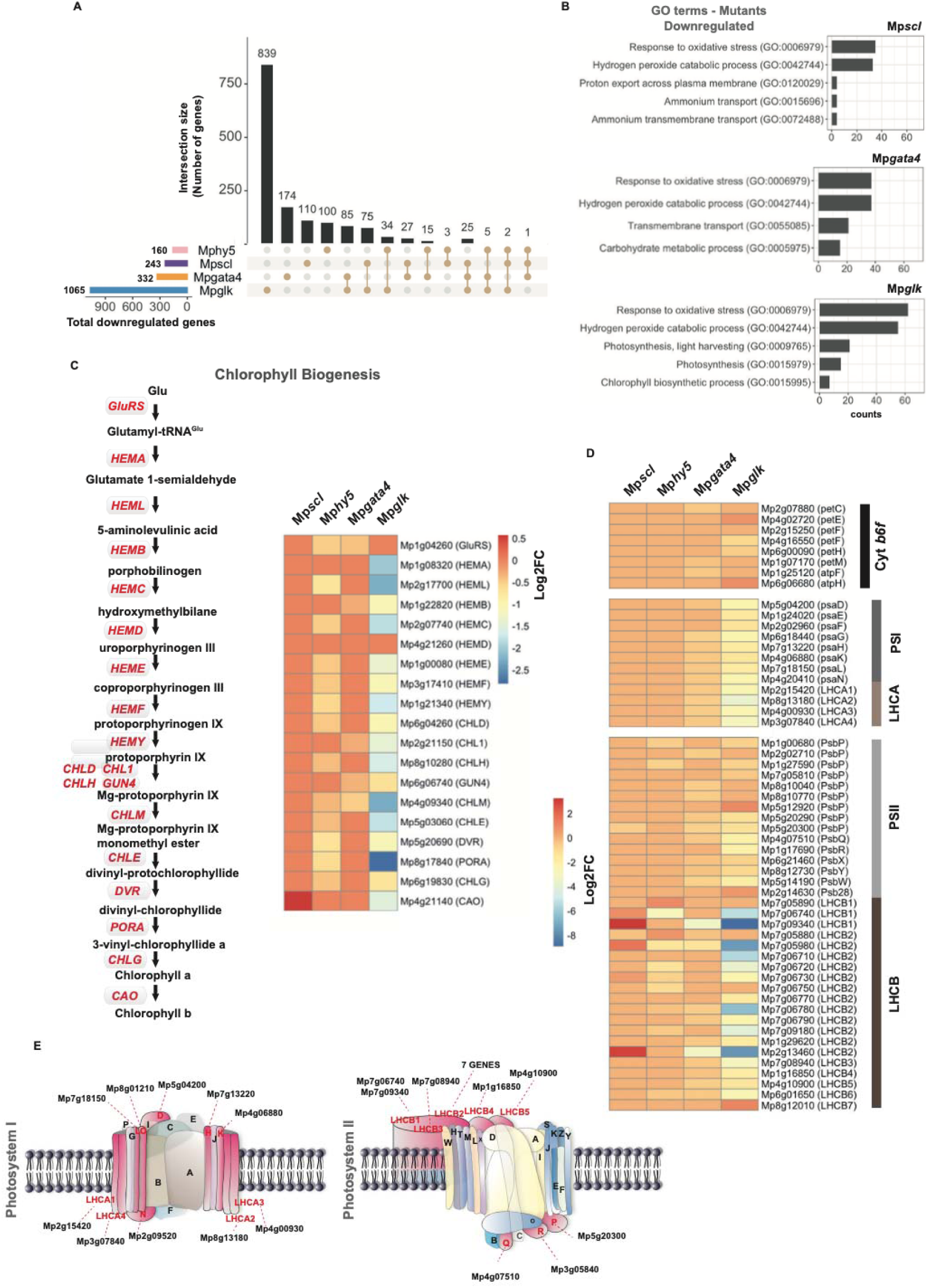
Mp*GLK* controls expression of photosynthesis associated genes. **A)** UpSet diagrams showing sets of downregulated genes in Mp*scl,* Mp*gata4,* Mp*hy5* and Mp*glk* mutants. **B)** Enriched Gene Ontology terms for Mp*scl,* Mp*gata4* and Mp*glk* mutants. **C)** Heatmap illustrating extent of down-regulation (Log2FoldChange) of transcripts encoding enzymes of chlorophyll biosynthesis in Mp*scl,* Mp*gata4,* Mp*hy5* and Mp*glk* mutant alleles. **D)** Heatmap indicating lower transcript abundance (Log2FoldChange) of genes encoding components of Photosystem II (PSII), Photosystem I (PSI), Light Harvest Complex A (LHCA), Light Harvest Complex B (LHCB) and cytochrome *b_6_f* complex in Mp*scl,* Mp*gata4,* Mp*hy5* and Mp*glk* mutants. **E)** Schematic representation of Photosystem I and II. Subunits showing an increase (>1) in corresponding transcript levels are highlighted in pink. Genes that are differentially expressed are highlighted in red. Figure modified from Water et al. 2009 and Tu et al. 2022.

We assessed the effects of Mp*scl*, Mp*gata4,* Mp*hy5* and Mp*glk* on the 68 annotated genes belonging to the light harvesting complexes A and B (LHCA and LHCB), components of photosystem I and II (PSI and PSII) and the cytochrome *b_6_f* complex. Mp*SCL* loss led to the upregulation of only two genes (Log2FC∼-0.37-0.55). Mp*HY5* or Mp*GATA4* loss resulted in a moderate reduction in transcript levels (Log2FC∼-0.4-0.7) of six and thirty genes respectively. The only exceptions were three and four *LHCB* genes where greater reductions (Log2FC>-1) were apparent for the Mp*gata4* and Mp*hy5* mutants respectively (**Figure 6D and Figure S7F**). Consistent with the effects on chlorophyll described above, the Mp*glk* mutant had the greatest effect on transcript abundance (**Figure 6D,E and Figure S7G**) with thirty-three of the sixty-eight genes being downregulated (Log2FC>=-1). HY5 is known to regulate cell elongation and proliferation, pigment accumulation as well as nutrient uptake^6^. When we examined whether homologs of genes known to control such processes in *A. thaliana* are mis-regulated in Mp*hy5* mutants, we only observed upregulation of a sulphate transporter (**Figure S7F**), a response opposite to that observed in *A. thaliana*. In conclusion, our RNA-SEQ analysis indicated that compared with MpGLK, MpSCL, MpHY5 and MpGATA4 have limited ability to control expression of photosynthesis genes, and their impact does not appear sufficiently extensive to impact on chloroplast biogenesis.

### Distinct and divergent binding patterns of MpGLK compared with flowering plants

To identify direct targets of MpGLK we performed ChIP-SEQ using transgenic *M. polymorpha* expressing MpGLK fused to a hemagglutinin (HA) tag (**Figure 7A**). Applying the data analysis pipeline utilised previously for five flowering plants (*A. thaliana*, tomato, rice, maize and tobacco)^40^ we detected a total of 2,654 reproducible peaks representing binding by MpGLK (IDR <0.01). Motif enrichment analysis of the summit regions of ChIP-SEQ peaks indicated MpGLK binds a RGATTYY sequence motif, analogous to its orthologs in flowering plants (**Figure 7B**). We next associated these peaks with 1.5 kb upstream regions of annotated genes and found 1,326 MpGLK targets (**Dataset S4**). We also performed assay for transposase-accessible chromatin with sequencing (ATAC-SEQ) to identify accessible chromatin regions and carried out ChIP-SEQ for H3K4me3, a histone mark often associated with active genes near their transcription start site. MpGLK ChIP signals overlapped with that from ATAC-SEQ (**Figure 7A, D**) suggesting that MpGLK binds to open chromatin. Binding sites were also found upstream of the +1 nucleosome often marked by H3K4me3, a pattern consistent with the observation for *A. thaliana* GLK (**Figure 7A, C**).

**Figure 7.**
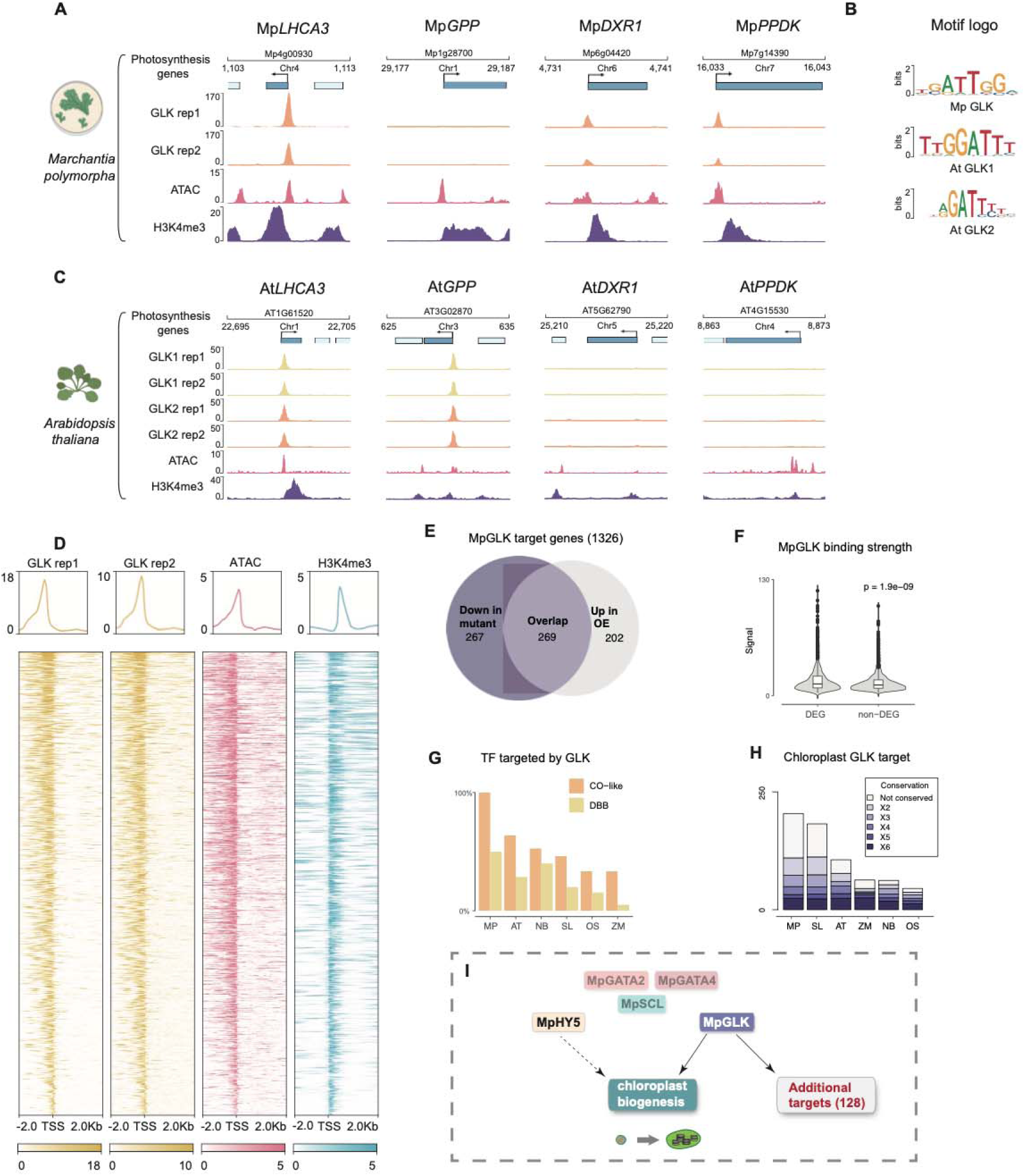
MpGLK ChIP-SEQ reveals divergence in GLK binding between *M. polymorpha* and flowering plants. **A)** Genome browser tracks showing GLK ChIP-SEQ peaks, open chromatin regions (ATAC-SEQ), and H3K4me3 peaks in conserved and *M. polymorpha* specific GLK target genes. *LHCB3: Light Harvest Complex 3; GPP: Galactose 1-phosphate phosphatase; DXR1; 1-deoxy-D-xylulose-5-phosphate reductoisomerase; PPDK: Pyruvate,orthophosphate (Pi) dikinase.* **B)** Motif enriched in the MpGLK and *A. thaliana* GLKs ChIP-SEQ peak regions. **C)** Genome browser tracks showing GLK1/2 binding, open chromatin and H3K4me3 in *A. thaliana* orthologs of *M. polymorpha* genes in panel A. **D)** Heatmap and average signal plot showing the chromatin features near the MpGLK binding sites. Clustering was performed using the GLK1 ChIP-SEQ rep1 signal. **E)** Partial overlap of MpGLK targets genes up-regulated in *GLK* over-expression (OE) lines and down-regulated in *GLK* knockout lines. **F)** Stronger ChIP-SEQ signals were found in differentially expressed GLK targets. **G)** Percentage of CONSTANS-Like (CO-like) and B-BOX (DBB) TF genes in each genome targeted by GLK. Note that all three CO-Like TFs in *M. polymorpha* are GLK targets. **H)** Number of predicted chloroplast localised GLK targets conserved between *M. polymorpha* and five flowering plants. **I)** Model summarising transcriptional regulation of chloroplast biogenesis in *M. polymorpha*.

Comparing ChIP-SEQ data with our gene expression data for the Mp*glk* mutants we found that 40.4% (536/1326) of GLK target genes were down-regulated in the Mp*glk* mutant (**Figure 7E**), and 35% (471/1326) were up-regulated in Mp*GLK* over-expression lines. Of those affected in Mp*glk* mutants and Mp*GLK* over-expression lines, 269 genes overlapped. As has been reported for *A. thaliana* and tomato, the average binding strength of MpGLK inferred from the ChIP-SEQ signal fold-change at the peak summit was higher in differentially expressed genes (DEGs) compared with non-DEGs (**Figure 7F**). Strikingly, more MpGLK target genes showed differential expression in Mp*glk* mutants compared with their counterparts in flowering plants. For example, only 21% (202/960) of AtGLK1/2 targets were differentially expressed in the At*glk1,glk2* double mutant. This is consistent with a more pronounced chlorophyll loss in Mp*glk* mutants described here compared with that for At*glk1,glk2* mutant alleles^11^. In summary, our findings suggest a conserved role for GLK throughout the evolution of land plants. It is possible of course that transcriptional networks in flowering plants have accumulated changes resulting in increased redundancy and robustness.

The dynamic nature of *cis*-regulatory elements in plant genomes can give rise to divergence in transcription factor binding and thus evolution of gene regulatory networks. For example, when GLK binding across flowering plants was compared only 10-20% of target genes were conserved between *A. thaliana,* tomato, tobacco, rice and maize. To investigate this for MpGLK, we classified protein-coding genes in *M. polymorpha* and the aforementioned five species into ortholog groups^41^. Only 53 GLK target genes in 48 ortholog groups were found to be conserved and they were mainly involved in photosynthesis (**Dataset S4**). For example, 23 genes encoding CHLOROPHYLL A/B BINDING PROTEIN were bound by MpGLK. Additionally, GLK bound CONSTANS and B-BOX transcription factor genes in all six species (**Figure 7G**) suggesting that they play an ancient regulatory role in circadian rhythm and photomorphogenesis. However, it is important to highlight that most MpGLK targets were not conserved. For instance, *1-deoxy-D-xylulose-5-phosphate reductoisomerase* and *Pyruvate,orthophosphate (Pi) dikinase* were bound by GLK in *M. polymorpha*, but their orthologs in *A. thaliana* were not bound by AtGLK1/2 (**Figure 7A and C**). Notably, even some MpGLK targets encoding chloroplast-localised proteins did not exhibit conservation between these species (**Figure 7H**). Our findings therefore provide strong support for the hypothesis that whilst GLK has maintained an important role in chloroplast biogenesis in land plants, the transcriptional network in which it operates has diverged significantly during the ∼400-500 MYA since the split between bryophyte and flowering plants.

## Discussion

### Loss of MpGATAs, MpHY5 or MpSCL has a minimal impact on chloroplast biogenesis

The monophyletic bryophyte group diverged from vascular plants approximately 400-500 MYA^21,23,25^. Since then, vascular plants and particularly angiosperms have undergone major morphological and physiological changes including elaborations in chloroplast biogenesis and photosynthesis. Very little is known about the evolution of the underlying genetic networks. We used *M. polymorpha* to test the extent to which regulators of chloroplast biogenesis defined in angiosperms are functionally conserved, and to shed light on the likely ancestral state.

Our analysis indicated that chlorophyll accumulation was not detectably perturbed in mutant alleles of Mp*gata4 or* Mp*gata2*. This contrasts with a ∼30-40% decrease in chlorophyll in *A. thaliana gnc*/*cga1* mutants^36^, and compromised photomorphogenesis when At*GATA2* expression was suppressed^19^. In *A. thaliana GNC* over-expression increased chlorophyll tenfold in seedlings and ∼30% in the leaf^15^, and *GATA2* over-expression led to constitutive photomorphogenesis^19^. The simplest and most parsimonious explanation for the lack of impact of loss of Mp*GATA4* or Mp*GATA2* function or over-expression on chlorophyll content in bryophytes such as *M. polymorpha* is that in the ancestral state these proteins do not play a role in chloroplast biogenesis. It is also possible that other proteins compensate and so redundancy in the gene regulatory networks masks loss of function. However, Mp*glk,gata4* and Mp*glk,gata2* double mutants had similar chlorophyll levels as Mp*glk* single mutants, implying lack of redundancy. It is also possible that GATA transcription factors regulate chloroplast biogenesis in some but not all bryophytes. This is perhaps supported by analysis of *P. patens* where two over-expression lines for Pp*GATA1*, one of the two *GNC/CGA1* orthologs in this species, showed a ∼10-20% increase in chlorophyll content^42^. A similar effect was reported when Pp*GATA1* was mis-expressed in *A. thaliana*^42^. However, these increases in chlorophyll content induced by Pp*GATA1* are modest compared with those reported in *A. thaliana* when *GNC* over-expression increased chlorophyll tenfold in seedlings and ∼30% in the leaf^15^. Moreover, while *GNC/CGA1* over-expression in *A. thaliana* led to an increase in chloroplast number per cell, this was not reported in *P. patens*^42^. The role of *PpGATA* orthologs (including both Pp*GATA1* and Pp*GATA2*), therefore needs additional analysis to confirm or refute their role in chloroplast biogenesis. A recent study in *M. polymorpha* proposed a role in response to high light stress^43^ for MpGATA4 with mutants showing mis-regulation of genes encoding EARLY LIGHT INDUCED PROTEINs (ELIPs). Our transcriptome analysis of Mp*gata4* mutants revealed that a single *ELIP* (Mp*ELIP-11*) was moderately downregulated (Log2FC=-0.678) alongside a small number of genes encoding components of the light harvesting complexes and photosystems. We also noted that editing Mp*GATA4* affected gametophyte morphology such that thalli of Mp*gata4* mutants had narrower lobes. This contrasts with *A. thaliana* where to our knowledge there have been no reports of perturbations to leaf morphology in *gnc*/*cga1* mutants. It is therefore possible that the role of this protein has been repurposed from one in affecting development of photosynthetic tissue and response to high light in *M. polymorpha*, to one in regulating chloroplast biogenesis in *A. thaliana*.

Similarly, neither Mp*scl* or Mp*mir171* mutants exhibited any detectable alterations in chlorophyll accumulation as would be expected if their functions were conserved between bryophytes and *A. thaliana.* For example, At*scl6* At*scl22* At*scl27* triple mutants and *MIR171* over-expressors lead to increased chlorophyll accumulation in *A. thaliana*^44^. As no other members of the *SCL* gene family in *M. polymorpha* contain a MIR171 recognition site, this argues against Mp*MIR171* playing a role in the regulation of chlorophyll accumulation. It is possible that the *M. polymorpha* homolog of At*SCL* does regulate chlorophyll biosynthesis but without the additional control derived from MIR171 found in *A. thaliana*. SCL6, SCL22 and SCL27 also regulate proliferation of meristematic cells^45–47^. However, if this were the case over-expression Mp*SCL* should decrease chlorophyll content as seen in *A. thaliana*^10^. However, our Mp*SCL* over-expression lines appeared similar to controls. A similar role in *M. polymorpha* could explain the perturbed thallus morphology with inward curling edges of Mp*scl* mutants. Thus, in addition to GATA2 and GATA4 discussed above, it is also possible that an ancestral role of SCL relates to development of photosynthetic tissue rather than chloroplast biogenesis *per se*. For SCL6, SCL22 and SCL27 this function seems to have been retained in *A. thaliana*, and a role in repressing chlorophyll synthesis acquired.

HY5 in *M. polymorpha* appears to play a conditional role in chloroplast biogenesis as has been reported in *A. thaliana*. Chlorophyll levels were comparable to controls when plants were grown at 22 or 17 °C. However, at 12 °C a small but statistically significant reduction in chlorophyll content was apparent in Mp*hy5*.

### A major and conserved role for GLK in chloroplast biogenesis in *M. polymorpha*

Unlike *GATAs* and the *MIR171-SCL* module, three lines of evidence indicate that GLK function is conserved in *M. polymorpha*. Firstly, Mp*glk* mutants have reduced chlorophyll accumulation and smaller chloroplasts with underdeveloped thylakoids. These perturbations to phenotype have been observed in other land plants^12–14,26,48^. Secondly, constitutive over-expression of Mp*GLK* resulted in increased chlorophyll accumulation, increased chloroplast size and ectopic chloroplast development similar to reports in angiosperms^36,49–52^. Thirdly, our transcriptome analysis of Mp*GLK* over-expression and mutant lines revealed an overlap between *GLK* dependent genes in *M. polymorpha* and *A. thaliana* but also rice, tobacco, tomato and maize^11,40^. ChIP-SEQ confirmed that most are photosynthesis-related genes directly bound by GLK such as the *LHCA*, *LHCB*, *PsbQ* and genes encoding chlorophyll biosynthesis enzymes. However, we also uncovered major divergence in GLK binding patterns associated with land plant evolution, as less than 5% of MpGLK target genes are conserved in flowering plants. Finally, and in contrast to previous reports, our phylogenetic analysis identified *GLK* orthologs in two Zygnematophyceae green algae for which genome assemblies became recently available^53^. Taken together our results suggest that the GLK function is ancestral to all land plants and appears to have evolved before the transition of plants to terrestrial ecosystems.

In summary, our results suggest that the regulation of photosynthesis gene expression is more streamlined in *M. polymorpha* (**Figure 7I**). It is possible that the low levels of genetic redundancy in regulators of photosynthesis in this species are associated with low rates of photosynthesis and limited specialisation within the thallus. Angiosperms on the other hand have a more complex development and morphology allowing colonisation of a broad range of environments. For example, leaves with specialised tissues allow high rates of photosynthesis. As a result, greater control over photosynthesis may have become necessary compared with bryophytes and could have been mediated by elaborations and specialisations to pre-existing pathways present in the common ancestor of bryophytes and vascular plants. The simplified genetic network in *M. polymorpha* either represents the ancestral state, or a simplified version due to secondary loss. Such secondary loss of traits has been reported for *M. polymorpha* stomata and the genetic network underpinning their development^54,55^. To elucidate the ancestral function of GLK further detailed genetic studies in other non-seed plants will be necessary. Furthermore, our finding of GLK homologs in the green algal lineage Zygnematophyceae, provides an exciting opportunity to test whether its role in photosynthesis predates the colonisation of land.

## Materials & Methods

### Phylogenetic analysis

To identify GATA B-Class and A-Class genes, three different approaches were combined: Firstly, the GATA protein sequences for 21 plant genomes were mined from the iTAK^56^ and PlantTFDB databases^57^, Phytozome, Fernbase, Phycozome and PhytoPlaza. Sequences for each individual species were aligned with the AtGNC and AtCGA1 amino acid sequences using MAFFT^58^. Results were filtered manually to identify GNC/CGA1 (B-Class) orthologs distinguished from other GATA family genes by the presence of conserved serine (S) residue, a conserved IRX(R/K)K motif (I: Isoleucine, R: Arginine, X: any amino acid and K: Lysine), and the presence or absence of conserved LLM-(leucine– leucine– methionine) domain at their C-terminus^31^. GATA2 (A-Class) orthologs, were distinguished by the presence of a conserved glutamine (Q) and a threonine (T) within the zinc finger motif. Secondly, we performed BLASTP searches against plant genomes on Phytozome v13, fern genomes (fernbase.org)^59^, hornwort genome (www.hornworts.uzh.ch)^21^, green algae genomes on PhycoCosm (phycocosm.jgi.doe.gov), and 1KP using the AtGNC/CGA1 amino acid sequence as a query. Results were filtered manually as described above. Finally, the combined results from the above to approaches were checked against Orthofinder searches^60^. The identified GATA protein sequences were aligned using MAFFT. Alignments were then trimmed using TrimAl^61^. A maximum likelihood phylogenetic tree was inferred using iQTree^62^, ModelFinder^63^ and ultrafast approximation for phylogenetic bootstrap^64^ and SH-aLRT test^65^. The tree was visualised using iTOL^66^.

To identify *GLK* genes, three different approaches were combined: Firstly, the G2-GARP protein sequences for 21 plant genomes were mined from the iTAK and PlantTFDB databases, Phytozome, Fernbase, Phycozome and PhytoPlaza. Sequences for each individual species were aligned with the AtGLK1/2 amino acid sequences using MAFFT. Results were filtered manually to identify GLK orthologs distinguished from other G2-GARP family genes by three characteristic motifs: AREAEAA motif (consensus motif) in the DNA-binding domain, VWG(Y/H)P and the PLGL(R/K)(P/S)P in the GCT-box domain. Secondly, we performed BLASTP searches against plant genomes on Phytozome v13, fern genomes (fernbase.org), hornworts genome (hwww.hornworts.uzh.ch), green algae genomes on PhycoCosm (/phycocosm.jgi.doe.gov), and 1KP^25^ using the AtGLK1/2 amino acid sequence as a query. Results were filtered manually as described above. Finally, the combined results from the above two approaches were checked against Orthofinder searches. The identified GLK protein sequences were aligned using MAFFT. Alignments were then trimmed using TrimAl. A maximum likelihood phylogenetic tree was inferred using iQTree, ModelFinder and ultrafast approximation for phylogenetic bootstrap and SH-aLRT test . The tree was visualised using iTOL .

### Plant growth, transformation, CRISPR/Cas9 gene editing and over-expression construct generation

*Marchantia polymorpha* accessions Cam-1 (male) and Cam-2 (female) were used in this study^67^ with the exception of ChIP-SEQ and ATAC-SEQ for which *Marchantia polymorpha* accessions Tak-1 (male) and Tak-2 (female) accessions were used. Plants were grown on half strength Gamborg B5 medium plus vitamins #G0210, Duchefa Biochemie) and 1.2% (w/v) agar (#A20021, Melford), under continuous light at 22 °C with light intensity of 100 μmol m^−2^ s^−1^. Transgenic *M. polymorpha* plants were obtained following an established protocol^28^.

For CRISPR/Cas9 gene editing, guide RNAs were *in silico* predicted using CasFinder tool (https://marchantia.info/tools/casfinder/). Several gRNAs per target gene were *in vitro* tested as described^68^ using oligonucleotides in **Dataset S5**. gRNA sequences that were selected to generate Mp*glk*, Mp*gata2*, Mp*gata4*, Mp*scl* and Mp*mir171* mutants are listed in **Dataset S5**. Single gRNA7 to mutate Mp*GLK* was cloned using OpenPlant toolkit^28^. Multiple gRNAs to mutate Mp*GATA4*, Mp*SCL* and Mp*MIR171* and a single gRNA to mutate Mp*GATA2* were cloned as described in^68^, using oligonucleotides listed in **Dataset S5** and the destination vector pMpGE013^69^. For the over-expression constructs, Mp*GLK and* Mp*GLKrw* CDS were synthesised (Integrated DNA Technologies), Mp*GLK* 3′UTR was amplified from *M. polymorpha* genomic DNA and cloned into the pUAP4 vector^28^. Constructs were generated using the OpenPlant toolkit^28^. OpenPlant parts used: OP-023 CDS12-eGFP, OP-020 CDS_hph, OP-037 CTAG_Lti6b, OP-054 3TER, _Nos-35S, OP-049 PROM_35S, OP-47 PROM_MpUBE2, OP-48 5UTR_MpUBE2, OP-063, L1_HyR-Ck1, OP073 L1_Cas9-Ck4, OP-076 L2_lacZgRNA-Cas9-CsA and L1_35S_s::eGFP-Lti6b^70^. Nucleotide sequence of Mp*GLKrw* CDS and oligonucleotide sequences used for cloning are listed in **Dataset S5**.

### Chlorophyll determination, fluorescence measurements and imaging analysis

For chlorophyll measurements, ∼30-50mg of 10-14 days old gemmalings were used, with five biological replicates per genotype. The tissue was blotted on tissue paper before weighing to remove excess water and then was transferred into a 1.5mL Eppendorf tube containing 1 mL of di-methyl sulfoxide (DMSO) (#D8418, Sigma Aldrich) and incubated in the dark at 65C with 300 rpm shaking for 45 minutes. Samples were let to cool down to room temperature for approximately one hour. Chlorophyll content was measured using a NanoDrop™One/One C Microvolume UV-Vis Spectrophotometer (ThermoFisher) following the manufacturer’s protocol. Chlorophyll fluorescence measurements were carried out using a CF imager (Technologica Ltd, UK) and the image processing software provided by the manufacturer, as described previously^71^. *M. polymorpha* plants were placed in the dark for 20 min for dark adaptation to evaluate the dark-adapted minimum fluorescence (Fo), dark-adapted maximum fluorescence (*Fm*), variable fluorescence *Fv* (*Fv*lJ*=*lJ*Fm–Fo*). All chlorophyll fluorescence images within each experiment were acquired at the same time in a single image, measuring a total of three plants per genotype and treatment. For the DCMU treatment, 20 µM DCMU (#45463, Sigma Aldrich) was added to the half-strength MS media before it was poured into the individual petri dishes. Thalli were placed for 24 h onto the DCMU-containing media before chlorophyll fluorescence measurements.

For imaging a gene frame (#AB0576, ThermoFisher) was positioned on a glass slide. Five to seven gemma were placed within the medium-filled gene frame together with 30 μL of milliQ water. The frame was then sealed with a cover slip. Plants were imaged immediately using a Leica SP8X spectral fluorescent confocal microscope. Imaging was conducted using either a 10× air objective (HC PL APO 10×/0.40 CS2) or a 20× air objective (HC PL APO 20×/0.75 CS2). Excitation laser wavelength and captured emitted fluorescence wavelength window were as follows: for eGFP (488 nm, 498−516 nm) and for chlorophyll autofluorescence (488 or 515nm, 670−700 nm). Chloroplast area was measured using ImageJ and the Macro in **Supplemental Information**.

For electron microscopy sections (∼2 mm^2^) of 5-6 individual 3-week-old *M. polymorpha* thallus per genotype were harvested; and then fixed, embedded and imaged using scanning electron microscopy as previously described^71^.

### Extraction of Terpenes and analysis on gas chromatograph - mass spectrometer

Approximately 200 mg of frozen *M. polymorpha* plant material was extracted with 1 mL of cold methanol containing 5 mM of NaCl to quench enzymatic activities^72^ and 900 ng of dodecane as internal standard (#297879, Sigma Aldrich). Mechanical disruption was achieved using a TissueLyser II (Qiagen), and the mixture was agitated on a Vibrax shaker at 2000 rpm for two hours. The samples were centrifuged to isolate the methanolic extracts devoid of plant debris, then extracted twice with hexane to capture non-polar and medium polar terpenes in the upper layer. Analysis of 200 μL hexane extracts was performed on a gas chromatograph (GC) Trace 1300 (ThermoFisher) coupled with a mass spectrometer (MS) ISQ 700 (ThermoFisher) and a CD-5MS column (30 m x 0.25 mm x 0.25 μm) (ThermoFisher).

Sample injection (1 μL) was conducted in splitless mode, following GC-MS oven parameters modified from^72^: initial temperature of 70°C for 3 min; first ramp of 20°C/min to 90°C; second ramp of 3°C/min to 180°C; third ramp of 5°C/min to 240°C; and final ramp of 20°C/min to 300°C, with a 6 min hold at this temperature. The MS initiated compound analysis after a 5.5-minute delay, with the transfer line and ion source temperatures set at 270°C and 230°C, respectively. Compounds were detected using the scan mode (scan time 0.17 s) within a mass detection range of 40 to 600 atomic mass units.

Chromatograms were processed using Chromeleon™ software (ThermoFisher), and quantification of main terpenes relied on comparison with the internal standard dodecane. Tentative identification of the major sesquiterpenes was achieved through comparison of mass spectra with literature, supplemented by a single sample run with a mix of C9-C40 alkanes standard to calculate the retention indices^73^.

### RNA extraction, cDNA preparation, qPCR **and** RNA sequencing

RNA was extracted from 3-4 two-week old gemmae, using the RNeasy Plant kit (#74903, Qiagen) according to the manufacturer’s protocol (RLT buffer supplemented with beta-merchaptoethanol was used) and residual genomic DNA was removed using the Turbo DNA-free kit (#AM1907, Invitrogen) according to the manufacturer’s instructions.

500 ng of DNase-treated RNA was used as a template for cDNA preparation using the SuperScript™ IV First-Strand Synthesis System (#18091050, Invitrogen) according to manufacturer’s instructions (with only modifying reverse transcriptase reaction time to 40 minutes and using oligo (dT)18 primers). qPCR was performed using the SYBR Green JumpStart *Taq* Ready Mix (#S4438, Sigma Aldrich) and a CFX384 RT System (Bio-Rad) thermal cycler. cDNA was diluted 6 times, oligonucleotides listed in **Dataset S5** were used at a final concentration of 0.5 μM and reaction conditions were as follows: initial denaturation step of 94°C for 2 minutes followed by 40 cycles of 94°C for 15 seconds (denaturation) and 60°C for 1 minute (annealing, extension, and fluorescence reading).

Library preparation and RNA sequencing was performed by Novogene (Cambridge, UK). Briefly, messenger RNA was purified from total RNA using poly-T oligo-attached magnetic beads. After fragmentation, the first strand cDNA was synthesised using random hexamer primers followed by the second strand cDNA synthesis. cDNA end repair, A-tailing, adapter ligation, size selection, amplification, and purification were performed next. Library concentration was measured on a Qubit instrument following the manufacturer’s procedure (ThermoFisher Scientific) followed by real-time qPCR quantification. Library size distribution was analysed on a bioanalyzer (Agilent) following the manufacturer’s protocol. Quantified libraries were pooled and sequenced on a NovaSeq PE150 Illumina platform and 6 G raw data per sample were obtained. Adapter sequences were: 5’ Adapter: 5’-AGATCGGAAGAGCGTCGTGTAGGGAAAGAGTGTAGATCTCGGTGGTCGCCGTATCATT-3’. 3’ Adapter: 5’-GATCGGAAGAGCACACGTCTGAACTCCAGTCACGGATGACTATCTCGTATGCCGTCTTCTGCTTG-3’

FastQC was used to assess read quality and TrimGalore (https://doi.org/10.5281/zenodo.5127899) to trim low-quality reads and remaining sequencing adapters. Reads were pseudo-aligned using Kallisto^74^ to the *M. polymorpha* Genome version 5 (primary transcripts only, obtained from MarpolBase)^75^. Kallisto estimates the abundance of each transcript in units of transcripts per million (TPM). Mapping statistics for each library are provided in **Dataset S3**. DGE analysis was performed with DESeq2^76^, with padj-values < 0.01, **Dataset S3**. Plots were generated using R. All raw data has been deposited in NCBI under the accession number PRJNA397637.

### ChIP-SEQ, ATAC-SEQ and data analysis

ChIP-SEQ was performed using two Mp*GLK* over-expression lines transformed with a construct of proMp*EF1a:*:Mp*GLK*-HA. Regenerating thalli from T1 transformants were cultured for 4 weeks and harvested at zeitgeber time 1h. Approximately 5g fresh thalli were crosslinked with 1% (w/v) formaldehyde. Nuclei were isolated as previously described^40^. Chromatin was then sonicated using Bioruptor (4x 5 min cycles of 30 sec ON/OFF). Sonicated chromatin was incubated with 2μg Anti-HA Tag Antibody (#26183, Invitrogen) for 6h at 4°C and an additional 2h at 4°C with 20μl Dynabeads Protein G (#10003D, ThermoFisher) blocked with 0.1% BSA. Beads with immunoprecipitated DNA were washed twice with low salt buffer (10mM Tris-HCl pH8.0, 0.15M NaCl, 1mM EDTA, 1% Triton X-100), twice with high salt buffer (10mM Tris-HCl pH8.0, 0.25M NaCl, 1mM EDTA, 1% Triton X-100) and TET (10mM Tris-HCl pH8.0, 1mM EDTA, 0.1% Tween-20). TS-Tn5 was purified and assembled with Tn5MEA/B adapter as previously described^77^. Tagmentation was performed on-bead with assembled TS-Tn5 for 30 min at 37°C. After tagmentation, beads were washed with low salt, high salt buffer and TET. Reverse crosslinking were performed at 55°C for 1h and 65°C overnight with 0.3M NaCl, 0.4% SDS, 10mM Tris-HCl pH8.0, 1mM EDTA, Proteinase K. DNA was purified with Qiagen MinElute (#28004, Qiagen) and PCR amplified with primers matching the Illumina N50x and N70x sequences. Libraries were sequenced with Hiseq X using the 150 bp paired-end mode. Histone ChIP-SEQ and ATAC-SEQ were performed as previously described^77^. All raw data has been deposited in NCBI under the accession number PRJNA1043823. The processed data can be viewed at http://www.epigenome.cuhk.edu.hk/.

ChIP-SEQ reads and ATAC-SEQ reads of *A. thaliana* and *M. polymorpha* were separately mapped to the reference genomes (*A. thaliana* TAIR10, *M. polymorpha* Tak v6) with bowtie2 with option -3 100 (version 2.3.5.1). The unmapped and low quality reads were filtered with SAMtools view (version 1.10) -F 4 and -q 20, and duplicated reads were removed using SAMTools rmdup. MACS2^78^ was used for peak calling and peak summit positions were retrieved by using the “–call-summits” function in MACS2. The peaks are resized to 150 bp regions (±75C:bp from the summit position) for IDR to calculate overlap. The summit’s signal fold-change values from two replicates are supplied to IDR to identify reproducible peaks. The overlap regions passed the IDR 0.01 and with average summit signal fold-change >5 in two replicates were kept and resized back to 150C:bp. They were then associated with gene promoters based on summit distance to the TSS (-1.5kb and + 0.5kb).

De novo motif discovery was performed with HOMER. GLK motifs in the GLK ChIP-SEQ summit region were extracted using findMotifsGenome.pl in HOMER, with the parameter - len 8. Orthofinder (version 2.2.7) was used to identify orthologs using the peptide sequence of all protein coding genes in *M. polymorpha*, *A. thaliana, Nicotiana benthamiana, Solanum lycopersicum, Oryza sativa,* and *Zea mays*. The transcription factor prediction was performed with iTAK. TargetP 2.0 (https://services.healthtech.dtu.dk/services/TargetP-2.0/) was used to predict chloroplast localization of the protein encoded by GLK target genes in Figure 7H. For example, the number of ortholog groups with GLK targets in 6 and 5 species were labelled as X6 and X5 respectively.

## Author contributions

N.Y.E., E.F., T.S., M.T., K. B., Ed.F., J.R., Z.M., S.R., Y.B., J.S-W., J.H. and S.Z. carried out the work. N.Y.E., E.F. and J.M.H. designed the work. N.Y.E, E.F. and J.M.H. wrote the manuscript with input from all authors.

## Acknowledgements

This work was funded as part of the BBSRC/EPSRC OpenPlant Synthetic Biology Research Centre Grant BB/ L014130/1 to J.H., BBSRC BB/F011458/1 for confocal microscopy to J.H, BBP0031171 to J.M.H. and Hong Kong GRF 1411918/14109420 to S.Z. T.B.S. was supported by the SNSF Postdoc Mobility Fellowship (P500PB_203128) and the EMBO Long-Term Fellowship (ALTF 531-2019). W.Z. was supported by the Zhujiang postdoc fellowship. J.S-W. was supported by the BBSRC DTP [BBSRC BB/X010899/1]. For the purpose of open access, the authors have applied a Creative Commons Attribution (CC BY) licence to any Author Accepted Manuscript version arising from this submission. We thank Karin H. Müller and Filomena Gallo from the Cambridge Advanced Imaging Centre for the electron microscopy sample preparation as well as the support during the image acquisition.

