## Supplementary material for "Streamlined regulation of chloroplast development in the liverwort *Marchantia polymorpha*": Supp Dataset 5

**SUPPLEMENTAL TABLES**

**Table 1. Oligonucleotides used in this study**

| **Name** | **Sequence** | **Description** |
| --- | --- | --- |
| MpGLKgRNA3-tr-F | GAAATTAATACGACTCACTATAGGGGGCGAGGGCTTCAAGATACGTTTTAGAGCTAGAAATAGC | *in vitro* gRNA testing |
| MpGLKgRNA7-tr-F | GAAATTAATACGACTCACTATAGGAGGATATGGAGTGGGTTGCTGTTTTAGAGCTAGAAATAGC | *in vitro* gRNA testing |
| MpGLKgRNA14-tr-F | GAAATTAATACGACTCACTATAGGTTGCAGCAGTGGAGAAAAGAGTTTTAGAGCTAGAAATAGC | *in vitro* gRNA testing |
| MpGLKgRNA17-tr-F | GAAATTAATACGACTCACTATAGGACCAATTGATCCAAATGTCTGTTTTAGAGCTAGAAATAGC | *in vitro* gRNA testing |
| GLK-Templ-F1 | GCAGAGTTCAAGTGGTGTAGC | *in vitro* gRNA testing, template amplification |
| GLK-Templ-R1 | TCCTGACGTAGCCTTGTTATTTAC | *in vitro* gRNA testing, template amplification |
| MpGLK-cDNA-R | GAGATCCTTACTTTATTTCCATTTGC | *in vitro* gRNA testing, template amplification, used with MpolyGLK1-UAP4-F (see below) |
| gRNA1-Mir171-tr-F | GAAATTAATACGACTCACTATAGGTGCTTACTGTGATGTTGGTGGTTTTAGAGCTAGAAATAGC | *in vitro* gRNA testing |
| gRNA2-Mir171-tr-F | GAAATTAATACGACTCACTATAGGGCGGCTGGGGACCAGGCATTGTTTTAGAGCTAGAAATAGC | *in vitro* gRNA testing |
| miR171-target-F | GCGCATCGATTCTGTCGTAAG | *in vitro* gRNA testing, template amplification |
| miR171-target-R | GGAACGAGCGTGAGCATG | *in vitro* gRNA testing, template amplification |
| gRNA1-MpScr-tr-F | GAAATTAATACGACTCACTATAGGTCGCGAGGGCCGACGGCATAGTTTTAGAGCTAGAAATAGC | *in vitro* gRNA testing |
| gRNA2-MpScr-tr-F | GAAATTAATACGACTCACTATAGGGCAGTCGAGCGTAGGGCACCGTTTTAGAGCTAGAAATAGC | *in vitro* gRNA testing |
| Mpoly_Scr_target-F | AGGCATGAATCACAATGAGGAAG | *in vitro* gRNA testing, template amplification |
| Mpoly_Scr_target-R | GGAATTGCACGAAGCTGTTG | *in vitro* gRNA testing, template amplification |
| MpGATA2-tr-g1 | GAAATTAATACGACTCACTATAGGAATCATATACCAACAAAGACGTTTTAGAGCTAGAAATAGC | *in vitro* gRNA testing |
| GATA2-templ-F2 | CCCGAGAGCAATTGACCCGA | *in vitro* gRNA testing, template amplification |
| GATA2-templ-R2 | GCTGTTCTCGTTGCCTTCGC | *in vitro* gRNA testing, template amplification |
| MpGATA4-tr-g1 | GAAATTAATACGACTCACTATAGGCTCTTTGTGGGCGTGGAGAAGTTTTAGAGCTAGAAATAGC | *in vitro* gRNA testing |
| MpGATA4-tr-g2 | GAAATTAATACGACTCACTATAGGACTACGACCACCGAGACAAGGTTTTAGAGCTAGAAATAGC | *in vitro* gRNA testing |
| MpGATA4_fw_templ | CTGACTGCACCCGAGACCTC | *in vitro* gRNA testing, template amplification |
| MpGATA4_rev_templ | TCCCAATGTCAGGAATTCCG | *in vitro* gRNA testing, template amplification |
| sg-in-vitro-R | AAAAGCACCGACTCGGTGCCAC | *in vitro* gRNA testing |
| L5AD5-J-F | CGGGTCTCAGGCAGGATGGGCAGTCTGCTCGAACAAAGCACCAGTGG | gRNA cloning, gRNA-tRNA scaffold |
| S5AD5-J-F | CGGGTCTCAGGCAGGATGGGCAGTCTGCTCG | gRNA cloning, gRNA-tRNA scaffold |
| S3AD5-J-R | TAGGTCTCCAAACGGATGAGCGACAGCAAACAAAAAAAAAAGCACCGACTCG | gRNA cloning, gRNA-tRNA scaffold |
| MpGLK-gRNA3-PTG-F | TAGGTCTCCGCTTCAAGATACGTTTTAGAGCTAGAA | gRNA cloning, gRNA-tRNA scaffold |
| MpGLK-gRNA3-PTG-R | ATGGTCTCAAAGCCCTCGCCCTGCACCAGCCGGGAA | gRNA cloning, gRNA-tRNA scaffold |
| MpGLK-gRNA7-PTG-F | TAGGTCTCCATGGAGTGGGTTGCTGTTTTAGAGCTAGAA | gRNA cloning, gRNA-tRNA scaffold |
| MpGLK-gRNA7-PTG-R | ATGGTCTCACCATATCCTTGCACCAGCCGGGAA | gRNA cloning, gRNA-tRNA scaffold |
| MpGLK-gRNA14-PTG-F | TAGGTCTCCGTGGAGAAAAGAGTTTTAGAGCTAGAA | gRNA cloning, gRNA-tRNA scaffold |
| MpGLK-gRNA14-PTG-R | ATGGTCTCACCACTGCTGCAATGCACCAGCCGGGAA | gRNA cloning, gRNA-tRNA scaffold |
| MpGLK-gRNA17-PTG-F | TAGGTCTCCTTGATCCAAATGTCTGTTTTAGAGCTAGAA | gRNA cloning, gRNA-tRNA scaffold |
| MpGLK-gRNA17-PTG-R | ATGGTCTCATCAATTGGTTGCACCAGCCGGGAA | gRNA cloning, gRNA-tRNA scaffold |
| MpGLK-gRNA-7-AarI-F | CTCGAGGATATGGAGTGGGTTGCT | Single gRNA cloning |
| MpGLK-gRNA-7-AarI-R | AAACAGCAACCCACTCCATATCCT | Single gRNA cloning |
| miR171-gRNA1-PTG-F | TAGGTCTCCTGTGATGTTGGTGGTTTTAGAGCTAGAA | gRNA cloning, gRNA-tRNA scaffold |
| miR171-gRNA1-PTG-R | ATGGTCTCACACAGTAAGCATGCACCAGCCGGGAA | gRNA cloning, gRNA-tRNA scaffold |
| miR171-gRNA2-PTG-F | TAGGTCTCCGGACCAGGCATTGTTTTAGAGCTAGAA | gRNA cloning, gRNA-tRNA scaffold |
| miR171-gRNA2-PTG-R | ATGGTCTCAGTCCCCAGCCGCTGCACCAGCCGGGAA | gRNA cloning, gRNA-tRNA scaffold |
| MpSCR-gRNA1-PTG-F | TAGGTCTCCGGCCGACGGCATAGTTTTAGAGCTAGAA | gRNA cloning, gRNA-tRNA scaffold |
| MpSCR-gRNA1-PTG-R | ATGGTCTCAGGCCCTCGCGATGCACCAGCCGGGAA | gRNA cloning, gRNA-tRNA scaffold |
| MpSCR-gRNA2-PTG-F | TAGGTCTCCAGCGTAGGGCACCGTTTTAGAGCTAGAA | gRNA cloning, gRNA-tRNA scaffold |
| MpSCR-gRNA2-PTG-R | ATGGTCTCACGCTCGACTGCTGCACCAGCCGGGAA | gRNA cloning, gRNA-tRNA scaffold |
| GATA2-gRNA1-PTG-F | TAGGTCTCCTATACCAACAAAGACGTTTTAGAGCTAGAA | gRNA cloning, gRNA-tRNA scaffold |
| GATA2-gRNA1-PTG-R | ATGGTCTCATATATGATTTGCACCAGCCGGGAA | gRNA cloning, gRNA-tRNA scaffold |
| MpGATA4-gRNA1-PTG-F | TAGGTCTCCGCGTGGAGAAGTTTTAGAGCTAGAA | gRNA cloning, gRNA-tRNA scaffold |
| MpGATA4-gRNA1-PTG-R | ATGGTCTCAACGCCCACAAAGAGTGCACCAGCCGGGAA | gRNA cloning, gRNA-tRNA scaffold |
| MpGATA4-gRNA2-PTG-F | TAGGTCTCCACCGAGACAAGGTTTTAGAGCTAGAA | gRNA cloning, gRNA-tRNA scaffold |
| MpGATA4-gRNA2-PTG-R | ATGGTCTCACGGTGGTCGTAGTTGCACCAGCCGGGAA | gRNA cloning, gRNA-tRNA scaffold |
| MpolyGLK1-UAP4-F | TTTGCTCTTCGTCTCAAATGTTTGCGTTCAAGGAGAAAT | Cloning Mp*GLK* CDS |
| MpolyGLK1-UAP4-R | TTTGCTCTTCGTCTCAAAGCCTACGAAGTGAGGGGAGGAG | Cloning Mp*GLK* CDS |
| MpGLK_rw_UAP4-F | TTTGCTCTTCGTCTCTAATGTTCGCCTTTAAGGAAAAGTTTC | Cloning Mp*GLKrw* CDS |
| MpGLK_rw_UAP4-R_stop | TTTGCTCTTCGTCTCTAAGCCTAGCTTGTCAGCGGTGGTG | Cloning Mp*GLKrw* CDS |
| OP-54_3TERM_Nos_35S_GGTA-F | TTTGCTCTTCGTCTCAGGTAGATCGTTCAAACATTTGGCA | Cloning Mp*GLK* 3´ UTR fused to Mp*GLK* CDS |
| OP-54_3TERM_Nos_35S_AGCG-R | TTTGCTCTTCGTCTCGAGCGATCTGGATTTTAGTACTGGATTTTG | Cloning Mp*GLK* 3´ UTR fused to Mp*GLK* CDS |
| MpGLK_3UTR-GCTT-F | TTTGCTCTTCGTCTCAGCTTGTATATAACGTAGTCATTGTTGTG | Cloning Mp*GLK* 3´ UTR fused to Mp*GLK* CDS |
| MpGLK_3UTR-TACC-R | TTTGCTCTTCGTCTCGTACCGAGATCCTTACTTTATTTCCATTTG | Cloning Mp*GLK* 3´ UTR fused to Mp*GLK* CDS |
| MpGLK3UTR-BsaF | TTTGGTCTCCGCTTCGTATATAACGTAGTCATTGTTGT | Cloning truncated versions of Mp*GLK* 3´ UTR fused to Mp*GLK* CDS |
| MpGLK3UTR-Bsa-100R | TTTGGTCTCCTACCGACAGCCTGCGCGATAAA | Cloning truncated versions of Mp*GLK* 3´ UTR fused to Mp*GLK* CDS |
| MpGLK3UTR-Bsa-100F | TTTGGTCTCCGCTTCATGAAACATGCTAAATCC | Cloning truncated versions of Mp*GLK* 3´ UTR fused to Mp*GLK* CDS |
| MpGLK3UTR-Bsa-200R | TTTGGTCTCCTACCCTTAGGCCCCAAAACAGC | Cloning truncated versions of Mp*GLK* 3´ UTR fused to Mp*GLK* CDS |
| MpGLK3UTR-Bsa-200F | TTTGGTCTCCGCTTAGCTATGAAGCTTTCTTCCT | Cloning truncated versions of Mp*GLK* 3´ UTR fused to Mp*GLK* CDS |
| MpGLK3UTR-Bsa-300R | TTTGGTCTCCTACCGTGGCTGTGGCTCTTTCA | Cloning truncated versions of Mp*GLK* 3´ UTR fused to Mp*GLK* CDS |
| MpGLK3UTR-Bsa-300F | TTTGGTCTCCGCTTTGCATAATATTTGCAGGC | Cloning truncated versions of Mp*GLK* 3´ UTR fused to Mp*GLK* CDS |
| MpGLK3UTR-Bsa-400R | TTTGGTCTCCTACCACTGAGTGGTTCTTGTTCTA | Cloning truncated versions of Mp*GLK* 3´ UTR fused to Mp*GLK* CDS |
| MpGLK3UTR-Bsa-400F | TTTGGTCTCCGCTTCGGGGAAGAGAAAAAGGA | Cloning truncated versions of Mp*GLK* 3´ UTR fused to Mp*GLK* CDS |
| MpGLK3UTR-Bsa-500R | TTTGGTCTCCTACCGCATTATAGTCTTGCCTATC | Cloning truncated versions of Mp*GLK* 3´ UTR fused to Mp*GLK* CDS |
| MpGLK3UTR-Bsa-500F | TTTGGTCTCCGCTTAAGTCTGCACGATCTTGTT | Cloning truncated versions of Mp*GLK* 3´ UTR fused to Mp*GLK* CDS |
| MpGLK3UTR-Bsa-600R | TTTGGTCTCCTACCTTGACAGACAGGTCAGGT | Cloning truncated versions of Mp*GLK* 3´ UTR fused to Mp*GLK* CDS |
| MpGLK3UTR-Bsa-600F | TTTGGTCTCCGCTTGTTAGTAACTTTGCCATGTCTAA | Cloning truncated versions of Mp*GLK* 3´ UTR fused to Mp*GLK* CDS |
| MpGLK3UTR-BsaR | TTTGGTCTCCTACCGAGATCCTTACTTTATTTCCATTTG | Cloning truncated versions of Mp*GLK* 3´ UTR fused to Mp*GLK* CDS |
| MpGLK-RT-F2 | TCCTACGATGGATCACACCG | qPCR |
| MpGLK-RT-R2 | CCCCAAGCATCAATACACGG | qPCR |
| MpGLK-rw-RT-F1 | TCTTCAGCTTGTCACCGAGT | qPCR |
| MpGLK-rw-RT-R1 | TGGCAATCGTTCGCATACAG | qPCR |
| MpGATA4-RT-F2 | GTAGCGAGGCGGACGAGAAC | qPCR |
| MpGATA4-RT-R2 | GTAGGAGCGGCGTCAGAAGT | qPCR |
| Mpgata2-qpcr-f1 | CGATCAGTACCACCGGCAGC | qPCR |
| Mpgata2-qpcr-r1 | CTCGTCACGGGTCTCGTTGC | qPCR |
| MpSCL-RT-F2 | CGCGCATGAAGATGGAGCAG | qPCR |
| MpSCL-RT-R2 | AGTAGCGCATGGTCTCGTGG | qPCR |
| MpACT_F | AGGCATCTGGTATCCACGAG | qPCR |
| MpACT_R | ACATGGTCGTTCCTCCAGAC | qPCR |
| MpAPT_F | CGAAAGCCCAAGAAGCTACC | qPCR |
| MpAPT_R | GTACCCCCGGTTGCAATAAG | qPCR |
| MpGLK-genot-gRNA1-Fw1 | AGACATGGCGTACGAAGGAT | Genotyping Mp*glk* CRISPR lines |
| MpGLK-genot-gRNA1-genot2 | TGGGAAGAGAAGACAGGGTGT | Genotyping Mp*glk* CRISPR lines |
| Olig-F1-genot | TGTAGTTTGAATGCAGGTTTGG | Genotyping Mp*glk* CRISPR lines |
| MpGLK_37_genot_F11 | TCTTATCAGTTACGATCGTAGGG | Genotyping Mp*glk* CRISPR lines |
| MpGLK_g37_genotR10 | CGAAACGGCGCAGAACATG | Genotyping Mp*glk* CRISPR lines |
| MpGATA4-genot-F | TGTACTGACAAGCGTTCGC | Genotyping Mp*gata4* CRISPR lines |
| MpGATA4-genot-R | TCTGATGGATCCACGACTG | Genotyping Mp*gata4* CRISPR lines |
| GATA2-genot-F1 | CTCGTTCGTGAGTGCGTGC | Genotyping Mp*ata2* CRISPR lines |
| GATA2-genot-R1 | GGGCACGGACAATTCTCGAT | Genotyping Mp*gata2* CRISPR lines |
| GATA2-genot-F2 | GGGGATGGATTCTTGCAGTTTGT | Genotyping Mp*gata2* CRISPR lines |
| GATA2-genot-R2 | AAAGTATGCGGGCTGTCGTG | Genotyping Mp*gata2* CRISPR lines |
| MpSCL_g234_CRISPR_F1 | AGGTGAGCGTGATGACTTCA | Genotyping Mp*scl* CRISPR lines |
| MpSCL_g234_CRISPR_R1 | ACCAGCTGCAATCCCGTC | Genotyping Mp*scl* CRISPR lines |
| 154-5-genot-F | GGTTGATGAAAGGCCGCAGTG | Genotyping Mp*mir171* CRISPR lines |
| 154-5-genot-R | CGTCTGCGCAGGATTCCGAA | Genotyping Mp*mir171* CRISPR lines |

**Table 2. Sequences of guide RNAs used in this study**

| **Target gene** | **Guide RNA sequence** |
| --- | --- |
| Mp*GLK*, gRNA3 | GGGCGAGGGCTTCAAGATAC |
| Mp*GLK*, gRNA7 | AGGATATGGAGTGGGTTGCT |
| Mp*GLK*, gRNA14 | TTGCAGCAGTGGAGAAAAGA |
| Mp*GLK*, gRNA17 | ACCAATTGATCCAAATGTCT |
| Mp*GATA4* gRNA*1* | CTCTTTGTGGGCGTGGAGAA |
| Mp*GATA4* gRNA*2* | ACTACGACCACCGAGACAAG |
| Mp*GATA2* gRNA1 | AATCATATACCAACAAAGAC |
| Mp*SCL* gRNA1 | TCGCGAGGGCCGACGGCATA |
| Mp*SCL* gRNA2 | GCAGTCGAGCGTAGGGCACC |
| Mp*MIR171* gRNA1 | TGCTTACTGTGATGTTGGTG |
| Mp*MIR171* gRNA2 | GCGGCTGGGGACCAGGCATT |
| Mp*HY5* gRNA*1* | caacttcagctcaaccgcaa |
| Mp*HY5* gRNA*2* | accagaactagaaagagggg |
