## Supplementary material for "Streamlined regulation of chloroplast development in the liverwort *Marchantia polymorpha*": File S1

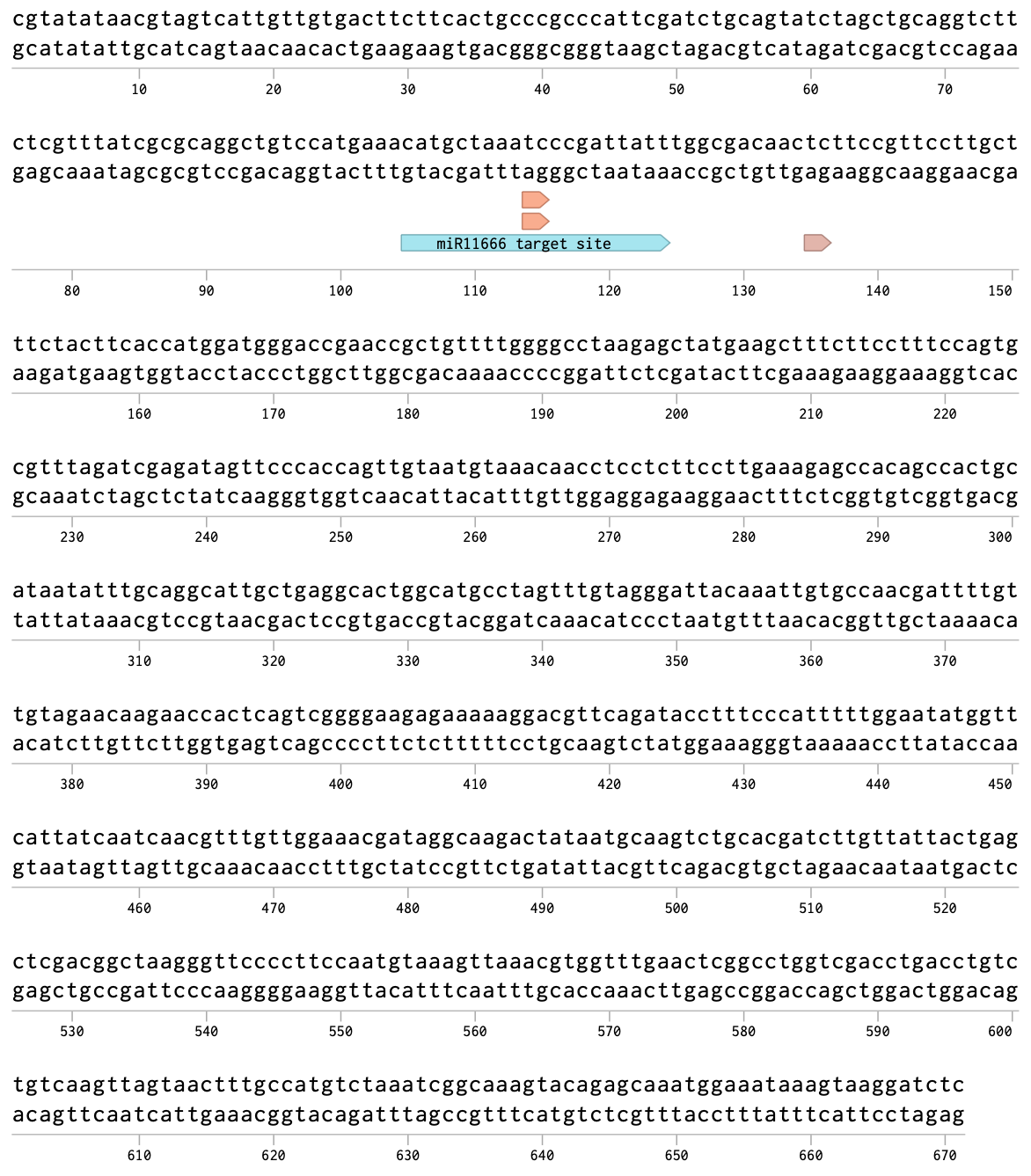

Mp*GLK 3´UTR miRNA and siRNA sites* (from Lin et al. 2016). with blue arrow indicating miR11666.4 binding site. Orange and light brown arrows indicating miR11666.4 and an unknown miRNA/siRNA putative cleavage sites, respectively.

```
atgttcgcctttaaggaaaagtttccggattggaaagactttccaaacggtcttagagtgttggtagttgatgaggatacgaaggtcctgaacgaaatcaagtgtaagcttgagggatgccaatatgttgttagcgcctttacaaagggtgaggacgctttggaggcacttcgggaccaacaaaatatattccatgtcgctctcgtcgaagctatgaccggggaagggtttaagattctcgaggtggctcgacaccttccaactattcttatgagtaacaccgagaatatggcgattatgatgcaaggtatcgcgttgggagccgctgacttcttgcagaagcctctctcagacgaaaagttgagaaatatatggcaacacgtagtgcggaaggccctgaatactggtgacccattgctcatggaaagtctggttccagtgaaggccgcagtcgagagtgtcttcagcttgtcaccgagtcctggacgggttaaggcggaaccagtatcttcccctggatcagacggcgaaagcaagagcaaagacatggaatgggtggccggcgttaccaagtcaccatcttcgggtgacgctgctgtatgcgaacgattgccagctccatctactcctcagcttgaacagactggccgcatgaatagccaggaggactcagtttcagaatctccaggctcagactcgttcgttcaggtcaaaacggaaggagaagaccccgagcctgacgcagctgccggtaaacctgcaccatgtactgaggtcaaggtggaagaccacgtcgaagcggtgtcagagtcatcacagctagatattaagttggaggactatccaacagttattaagttagagttagacgacaacatagacgacattggtctgtcgaacgggatcgtcgttgacggagacggaggctcaggtctagatattaacccatgtctgttagttccactgccggaaagtgcactcgacatgcatatgaacatcgaagcgcttgacgctggtgaggtgcatgatacagaaatcggcgaggaagaagcgcttcttttggctgaggttgctaaagcaggtgctgaggggctggagcggacgccatcaatgaactgtttgtcggtggattccggtgactgttcgtccggtgagaaaaaaggagaaggatgtaaggttaacaataaagcgacctctggtcgacgcaagatgaaagtggattggacgccagaactccacagaagattcgttcaggctgttgagcagctgggtgtggaaaaagccataccttcaaggatcctagaactgatgggtgttcagtgcttaacaaggcataacatcgcttcgcacctccaaaagtatcggtcacaccggcgccacttgcttgctcgagaggcggaggcggccacatggcaccaccgtaagcctatagaccctaacgtttgggcccgctcccgaagagacggaacagcttggctcgcaccacaccacaccaatccacctcctatccagccaaggcctcctatgggtctgaccccaattcagccccaccctggagcgcattgccatccgatgggtccccccatgcacgtgtggggacacccaaccatggaccatacggcggctcatatgtggcaacaacctcagatggcaactcctactacttggcaggccccagacggtagctactggcaacacccctgcatcgacgcgtggggacaccctactccaggacccggtacaccctgttaccctcagccttatcgagtgcctatggcgccgatgcccgcttttgcttctccgatgactacagcggctttggccgcagattcgtactttgcagacgaaagcatgccaattccgatgtatccgacagctccggacgatcccgagcttacagttgctgcgggtgccgctgcgtcgtcgaagccatctgattttcatccccctaaagaaatcctcgatgcagccatttccgaggccttggctaatccctggactccccttcctcttggacttaagccaccgtcaatggaaggagtcatggcggaactccagagacagggtattaatactgtacctccaccaccgctgacaagc

Nucleotide sequence alignment
wt_MpGLK             atgtttgcgttcaaggagaaattccccgactggaaggatttccctaatggattgcgggtt       60
rewritten_MpGLK      atgttcgcctttaaggaaaagtttccggattggaaagactttccaaacggtcttagagtg       60
                     ***** ** ** ***** ** ** ** ** ***** ** ** ** ** **  *  * **

wt_MpGLK             cttgttgtcgacgaagacaccaaagtgctcaatgagattaagtgcaagttggaaggttgt       120
rewritten_MpGLK      ttggtagttgatgaggatacgaaggtcctgaacgaaatcaagtgtaagcttgagggatgc       120
                      * ** ** ** ** ** ** ** ** ** ** ** ** ***** *** * ** ** **

wt_MpGLK             cagtatgtcgtctcggctttcactaaaggagaagatgcccttgaagcattgcgagatcag       180
rewritten_MpGLK      caatatgttgttagcgcctttacaaagggtgaggacgctttggaggcacttcgggaccaa       180
                     ** ***** **    ** ** ** ** ** ** ** **  * ** *** * ** ** **

wt_MpGLK             cagaacatttttcacgtggcgctggtggaggcaatgacgggcgagggcttcaagatactg       240
rewritten_MpGLK      caaaatatattccatgtcgctctcgtcgaagctatgaccggggaagggtttaagattctc       240
                     ** ** ** ** ** ** ** ** ** ** ** ***** ** ** ** ** ***** **

wt_MpGLK             gaagttgcaagacatttgcctacaatattgatgtccaatacggaaaacatggccatcatg       300
rewritten_MpGLK      gaggtggctcgacaccttccaactattcttatgagtaacaccgagaatatggcgattatg       300
                     ** ** **  ****  * ** ** **  * ***   ** ** ** ** ***** ** ***

wt_MpGLK             atgcagggaattgcgcttggtgcagcggattttcttcaaaagccactgtctgatgagaag       360
rewritten_MpGLK      atgcaaggtatcgcgttgggagccgctgacttcttgcagaagcctctctcagacgaaaag       360
                     ***** ** ** *** * ** ** ** ** **  * ** ***** ** ** ** ** ***

wt_MpGLK             cttcggaacatttggcagcatgttgttcgaaaggctctcaacacaggtgatcctcttctg       420
rewritten_MpGLK      ttgagaaatatatggcaacacgtagtgcggaaggccctgaatactggtgacccattgctc       420
                      *  * ** ** ***** ** ** ** ** ***** ** ** ** ***** **  * **

wt_MpGLK             atggagtccctcgtacctgttaaagcagctgtggaatccgtgttttcgctttctccctcc       480
rewritten_MpGLK      atggaaagtctggttccagtgaaggccgcagtcgagagtgtcttcagcttgtcaccgagt       480
                     *****    ** ** ** ** ** ** ** ** **    ** **     * ** **

wt_MpGLK             ccaggtcgagtcaaagctgagcctgtttcaagtccaggttctgatggggagtcgaagtcg       540
rewritten_MpGLK      cctggacgggttaaggcggaaccagtatcttcccctggatcagacggcgaaagcaagagc       540
                     ** ** ** ** ** ** ** ** ** **    ** ** ** ** ** **    ***

wt_MpGLK             aaggatatggagtgggttgctggggtgacgaaatctccttcaagcggagatgcagcagtt       600
rewritten_MpGLK      aaagacatggaatgggtggccggcgttaccaagtcaccatcttcgggtgacgctgctgta       600
                     ** ** ***** ***** ** ** ** ** ** ** ** **    ** ** ** ** **

wt_MpGLK             tgtgagagacttcctgcaccttcaacaccacagttggagcaaacagggaggatgaactcg       660
rewritten_MpGLK      tgcgaacgattgccagctccatctactcctcagcttgaacagactggccgcatgaatagc       660
                     ** **  ** * ** ** ** ** ** ** *** * ** ** ** **  * *****

wt_MpGLK             caagaagattcagtgtcagagtcacctgggtctgatagctttgtgcaagtgaagaccgag       720
rewritten_MpGLK      caggaggactcagtttcagaatctccaggctcagactcgttcgttcaggtcaaaacggaa       720
                     ** ** ** ***** ***** ** ** ** ** **    ** ** ** ** ** ** **

wt_MpGLK             ggtgaggatccggaaccagatgctgccgcaggaaagccagcaccttgcacagaagtgaaa       780
rewritten_MpGLK      ggagaagaccccgagcctgacgcagctgccggtaaacctgcaccatgtactgaggtcaag       780
                     ** ** ** ** ** ** ** ** ** ** ** ** ** ***** ** ** ** ** **

wt_MpGLK             gttgaggatcatgtggaggctgtttctgaatcttctcaattagacatcaagcttgaagat       840
rewritten_MpGLK      gtggaagaccacgtcgaagcggtgtcagagtcatcacagctagatattaagttggaggac       840
                     ** ** ** ** ** ** ** ** ** ** ** ** **  **** ** *** * ** **

wt_MpGLK             tatcctactgtcatcaagctagaattagatgataatatagatgatatcggactcagcaat       900
rewritten_MpGLK      tatccaacagttattaagttagagttagacgacaacatagacgacattggtctgtcgaac       900
                     ***** ** ** ** *** **** ***** ** ** ***** ** ** ** **    **

wt_MpGLK             ggcattgtggtagatggtgatggtgggtctggattagacatcaatccttgcctcctagta       960
rewritten_MpGLK      gggatcgtcgttgacggagacggaggctcaggtctagatattaacccatgtctgttagtt       960
                     ** ** ** ** ** ** ** ** ** ** **  **** ** ** ** ** **  ****

wt_MpGLK             cctctccccgagtccgctctggatatgcacatgaatattgaggctttggatgccggagaa       1020
rewritten_MpGLK      ccactgccggaaagtgcactcgacatgcatatgaacatcgaagcgcttgacgctggtgag       1020
                     ** ** ** **    ** ** ** ***** ***** ** ** **  * ** ** ** **

wt_MpGLK             gttcacgacactgagattggggaagaggaggcccttttgcttgcagaagtcgcgaaggct       1080
rewritten_MpGLK      gtgcatgatacagaaatcggcgaggaagaagcgcttcttttggctgaggttgctaaagca       1080
                     ** ** ** ** ** ** ** ** ** ** ** *** *  * ** ** ** ** ** **

wt_MpGLK             ggagccgaaggcctcgaacgaaccccttctatgaattgccttagcgttgacagtggagat       1140
rewritten_MpGLK      ggtgctgaggggctggagcggacgccatcaatgaactgtttgtcggtggattccggtgac       1140
                     ** ** ** ** ** ** ** ** ** ** ***** **  *    ** **    ** **

wt_MpGLK             tgcagcagtggagaaaagaagggtgagggttgcaaagtaaataacaaggctacgtcagga       1200
rewritten_MpGLK      tgttcgtccggtgagaaaaaaggagaaggatgtaaggttaacaataaagcgacctctggt       1200
                     **       ** ** ** ** ** ** ** ** ** ** ** ** ** ** ** ** **

wt_MpGLK             agaaggaaaatgaaggttgactggacccctgagctgcatcggcggtttgtccaggcagtc       1260
rewritten_MpGLK      cgacgcaagatgaaagtggattggacgccagaactccacagaagattcgttcaggctgtt       1260
                      ** * ** ***** ** ** ***** ** ** ** **  *  * ** ** ***** **

wt_MpGLK             gaacaactcggagttgagaaggctattccatctcgcattttagagctcatgggagtacag       1320
rewritten_MpGLK      gagcagctgggtgtggaaaaagccataccttcaaggatcctagaactgatgggtgttcag       1320
                     ** ** ** ** ** ** ** ** ** ** **  * **  **** ** ***** ** ***

wt_MpGLK             tgtctaactcgtcacaatattgccagccatctgcagaaatatcgatctcatcgaaggcat       1380
rewritten_MpGLK      tgcttaacaaggcataacatcgcttcgcacctccaaaagtatcggtcacaccggcgccac       1380
                     **  ****  * ** ** ** **    ** ** ** ** ***** ** ** **  * **

wt_MpGLK             cttttggcgagagaagctgaagccgcgacttggcatcatcgcaaaccaattgatccaaat       1440
rewritten_MpGLK      ttgcttgctcgagaggcggaggcggccacatggcaccaccgtaagcctatagaccctaac       1440
                      *  * **  **** ** ** ** ** ** ***** ** ** ** ** ** ** ** **

wt_MpGLK             gtctgggctaggagtagacgggatggtactgcgtggctggcccctcatcatacgaaccct       1500
rewritten_MpGLK      gtttgggcccgctcccgaagagacggaacagcttggctcgcaccacaccacaccaatcca       1500
                     ** *****  *     ** * ** ** ** ** ***** ** ** ** ** ** ** **

wt_MpGLK             ccaccaattcagcctcgcccaccaatgggactcacgcctatacagccgcatccaggtgct       1560
rewritten_MpGLK      cctcctatccagccaaggcctcctatgggtctgaccccaattcagccccaccctggagcg       1560
                     ** ** ** *****  * ** ** ***** ** ** ** ** ***** ** ** ** **

wt_MpGLK             cactgtcaccccatgggaccgccgatgcatgtttggggtcatcctacgatggatcacacc       1620
rewritten_MpGLK      cattgccatccgatgggtccccccatgcacgtgtggggacacccaaccatggaccatacg       1620
                     ** ** ** ** ***** ** ** ***** ** ***** ** ** ** ***** ** **

wt_MpGLK             gctgcgcacatgtggcagcagccacaaatggccacaccaacaacatggcaagcacctgat       1680
rewritten_MpGLK      gcggctcatatgtggcaacaacctcagatggcaactcctactacttggcaggccccagac       1680
                     ** ** ** ******** ** ** ** ***** ** ** ** ** ***** ** ** **

wt_MpGLK             ggatcgtattggcagcatccgtgtattgatgcttggggtcatccaacacctggtccggga       1740
rewritten_MpGLK      ggtagctactggcaacacccctgcatcgacgcgtggggacaccctactccaggacccggt       1740
                     **    ** ***** ** ** ** ** ** ** ***** ** ** ** ** ** ** **

wt_MpGLK             actccgtgctatccacagccatacagagttccaatggctcctatgccggcgttcgcgtca       1800
rewritten_MpGLK      acaccctgttaccctcagccttatcgagtgcctatggcgccgatgcccgcttttgcttct       1800
                     ** ** ** ** ** ***** **  **** ** ***** ** ***** ** ** ** **

wt_MpGLK             cccatgacaactgctgcccttgcagccgacagctatttcgctgatgagtcgatgcctata       1860
rewritten_MpGLK      ccgatgactacagcggctttggccgcagattcgtactttgcagacgaaagcatgccaatt       1860
                     ** ***** ** ** **  * ** ** **    ** ** ** ** **    ***** **

wt_MpGLK             cccatgtaccccactgcacccgatgacccggaattgactgtcgcggctggcgctgcggcc       1920
rewritten_MpGLK      ccgatgtatccgacagctccggacgatcccgagcttacagttgctgcgggtgccgctgcg       1920
                     ** ***** ** ** ** ** ** ** ** **  * ** ** ** ** ** ** ** **

wt_MpGLK             agcagcaagccttcagacttccacccgccaaaggagattctggacgccgcaatcagtgaa       1980
rewritten_MpGLK      tcgtcgaagccatctgattttcatccccctaaagaaatcctcgatgcagccatttccgag       1980
                           ***** ** ** ** ** ** ** ** ** ** ** ** ** ** **    **

wt_MpGLK             gctttggccaacccgtggacaccgttgccacttggtttgaagcctccctctatggagggt       2040
rewritten_MpGLK      gccttggctaatccctggactccccttcctcttggacttaagccaccgtcaatggaagga       2040
                     ** ***** ** ** ***** **  * ** *****  * ***** ** ** ***** **

wt_MpGLK             gtgatggctgagctccagcggcagggaatcaacacagttccacctcctcccctcacttcg       2100
rewritten_MpGLK      gtcatggcggaactccagagacagggtattaatactgtacctccaccaccgctgacaagc       2100
                     ** ***** ** ****** * ***** ** ** ** ** ** ** ** ** ** **

wt_MpGLK             tag  2103
rewritten_MpGLK      ---  2100
```

Translation Mp*GLK* re-written:

MFAFKEKFPDWKDFPNGLRVLVVDEDTKVLNEIKCKLEGCQYVVSAFTKGEDALEALRDQQNIFHVALVEAMTGEGFKILEVARHLPTILMSNTENMAIMMQGIALGAADFLQKPLSDEKLRNIWQHVVRKALNTGDPLLMESLVPVKAAVESVFSLSPSPGRVKAEPVSSPGSDGESKSKDMEWVAGVTKSPSSGDAAVCERLPAPSTPQLEQTGRMNSQEDSVSESPGSDSFVQVKTEGEDPEPDAAAGKPAPCTEVKVEDHVEAVSESSQLDIKLEDYPTVIKLELDDNIDDIGLSNGIVVDGDGGSGLDINPCLLVPLPESALDMHMNIEALDAGEVHDTEIGEEEALLLAEVAKAGAEGLERTPSMNCLSVDSGDCSSGEKKGEGCKVNNKATSGRRKMKVDWTPELHRRFVQAVEQLGVEKAIPSRILELMGVQCLTRHNIASHLQKYRSHRRHLLAREAEAATWHHRKPIDPNVWARSRRDGTAWLAPHHTNPPPIQPRPPMGLTPIQPHPGAHCHPMGPPMHVWGHPTMDHTAAHMWQQPQMATPTTWQAPDGSYWQHPCIDAWGHPTPGPGTPCYPQPYRVPMAPMPAFASPMTTAALAADSYFADESMPIPMYPTAPDDPELTVAAGAAASSKPSDFHPPKEILDAAISEALANPWTPLPLGLKPPSMEGVMAELQRQGINTVPPPPLTS

Translation wt Mp*GLK*:

MFAFKEKFPDWKDFPNGLRVLVVDEDTKVLNEIKCKLEGCQYVVSAFTKGEDALEALRDQQNIFHVALVEAMTGEGFKILEVARHLPTILMSNTENMAIMMQGIALGAADFLQKPLSDEKLRNIWQHVVRKALNTGDPLLMESLVPVKAAVESVFSLSPSPGRVKAEPVSSPGSDGESKSKDMEWVAGVTKSPSSGDAAVCERLPAPSTPQLEQTGRMNSQEDSVSESPGSDSFVQVKTEGEDPEPDAAAGKPAPCTEVKVEDHVEAVSESSQLDIKLEDYPTVIKLELDDNIDDIGLSNGIVVDGDGGSGLDINPCLLVPLPESALDMHMNIEALDAGEVHDTEIGEEEALLLAEVAKAGAEGLERTPSMNCLSVDSGDCSSGEKKGEGCKVNNKATSGRRKMKVDWTPELHRRFVQAVEQLGVEKAIPSRILELMGVQCLTRHNIASHLQKYRSHRRHLLAREAEAATWHHRKPIDPNVWARSRRDGTAWLAPHHTNPPPIQPRPPMGLTPIQPHPGAHCHPMGPPMHVWGHPTMDHTAAHMWQQPQMATPTTWQAPDGSYWQHPCIDAWGHPTPGPGTPCYPQPYRVPMAPMPAFASPMTTAALAADSYFADESMPIPMYPTAPDDPELTVAAGAAASSKPSDFHPPKEILDAAISEALANPWTPLPLGLKPPSMEGVMAELQRQGINTVPPPPLTS

**MpGLK*-CDSrw* nucleotide and protein sequence.** Mp*GLK* CDS “re-written” nucleotide sequence, its alignment to Mp*GLK* native CDS sequence and translation of both Mp*GLK* ‘re-written’ and native open reading frames.
