## Supplementary material for "Streamlined regulation of chloroplast development in the liverwort *Marchantia polymorpha*": Supp Info

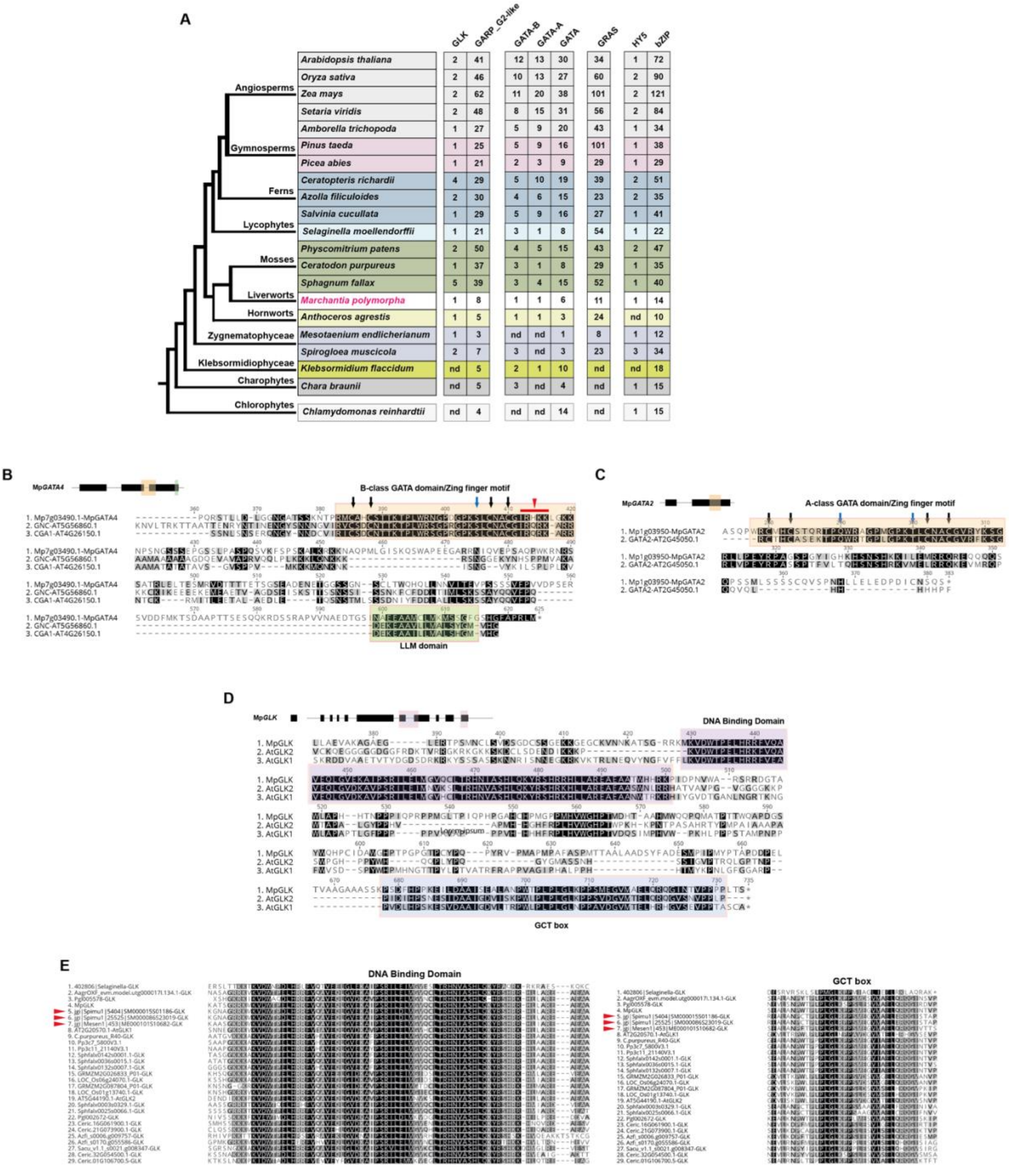


**Figure S1. Phylogenetic analysis of GARP, GATA, GRAS and bZIP transcription factors. A)** Phylogenetic relationships of the major lineages of land plants and green algae (Li et al. 2020). Gene numbers (indicated with numbered squares, nd= non-detected) of GARP G2-like, GATA and GRAS transcription factor families and subfamilies, for representative species for which high quality genome assemblies are available. **B)** MAFFT alignment of MpGATA4 and *A. thaliana* GNC and CGA1 proteins. The B-Class GATA domain is highlighted with orange and the LLM domain is highlighted with green, both in the amino acid alignment and the gene model on the top of the alignment. Conserved cysteines within the zinc finger motif are indicated with black arrows. The conserved S residue is indicated with a blue arrow and conserved IRX(R/K)K sequence is indicated with a red arrowhead and bar. **C)** MAFFT alignment of MpGATA2 and the *A. thaliana* GATA2 protein*.* A-Class GATA domain is highlighted with orange, both in the amino acid alignment and the gene model on the top of the alignment. The conserved cysteines within the zinc finger motif are indicated with black arrows. The conserved Q and T are indicated with blue arrows. The colouring used for that column depends on the fraction of the column that is made of letters from this group. Black: 100% similar, dark-gray: 80−100% similar, lighter gray: 60−80% similar, white: less than 60% similar. **D)** MAFFT alignment of MpGLK and AtGLK1 and 2 proteins*.* The GLK DNA binding domain is highlighted with purple and the GCT box is highlighted with blue, both in the amino acid alignment and the gene model on the top of the alignment. The colouring used for that column depends on the fraction of the column that is made of letters from this group. Black: 100% similar, dark-gray: 80−100% similar, lighter gray: 60−80% similar, white: less than 60% similar. **E)** Amino acid sequence alignment of different GLK proteins. Red arrowheads indicate green algae sequences.

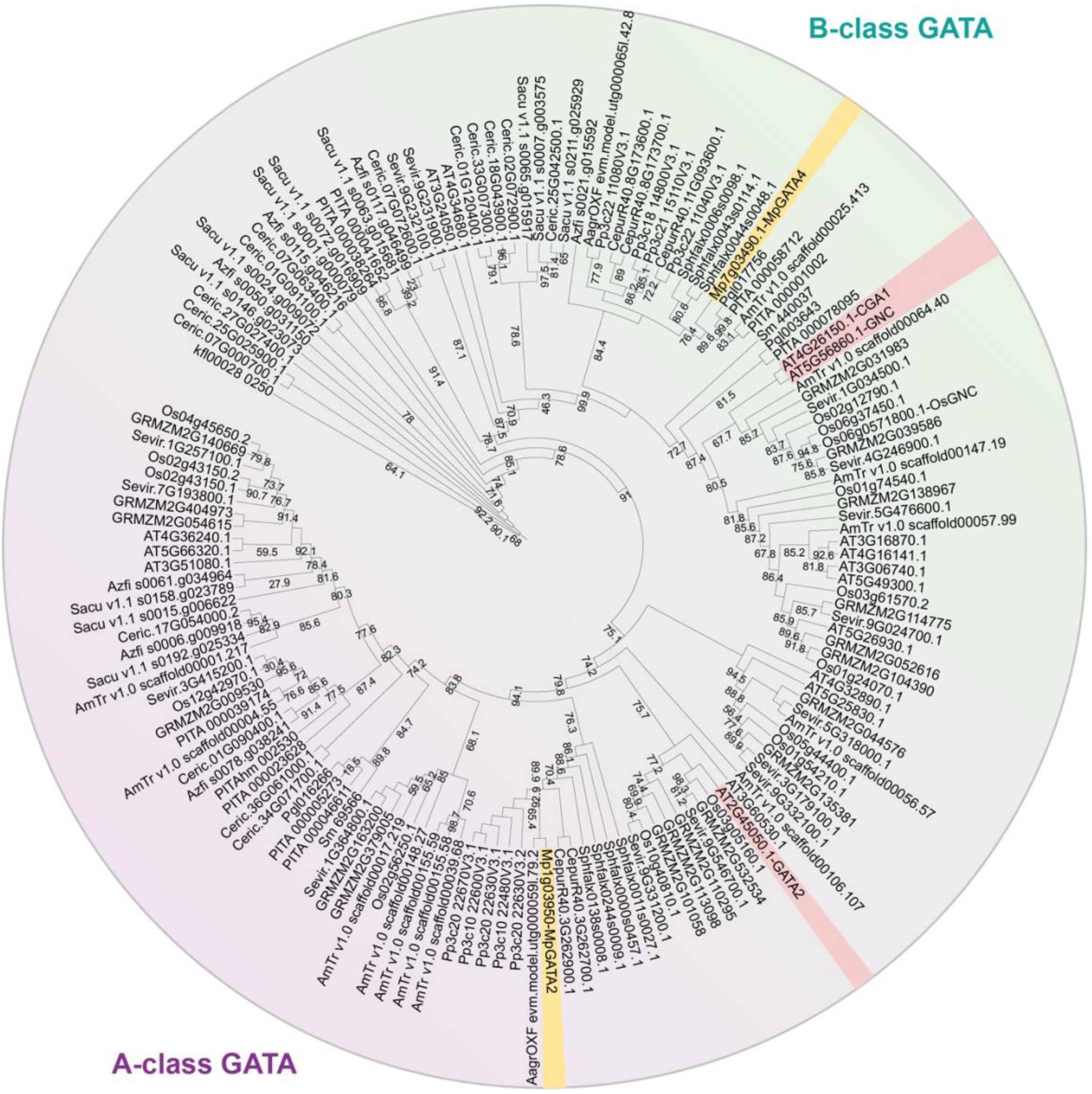


**Figure S2. Phylogenetic analysis of GATA transcription factors.**
GATA phylogeny. Numbers on branches represent SH-aLRT test support (Guindon et al. 2010).

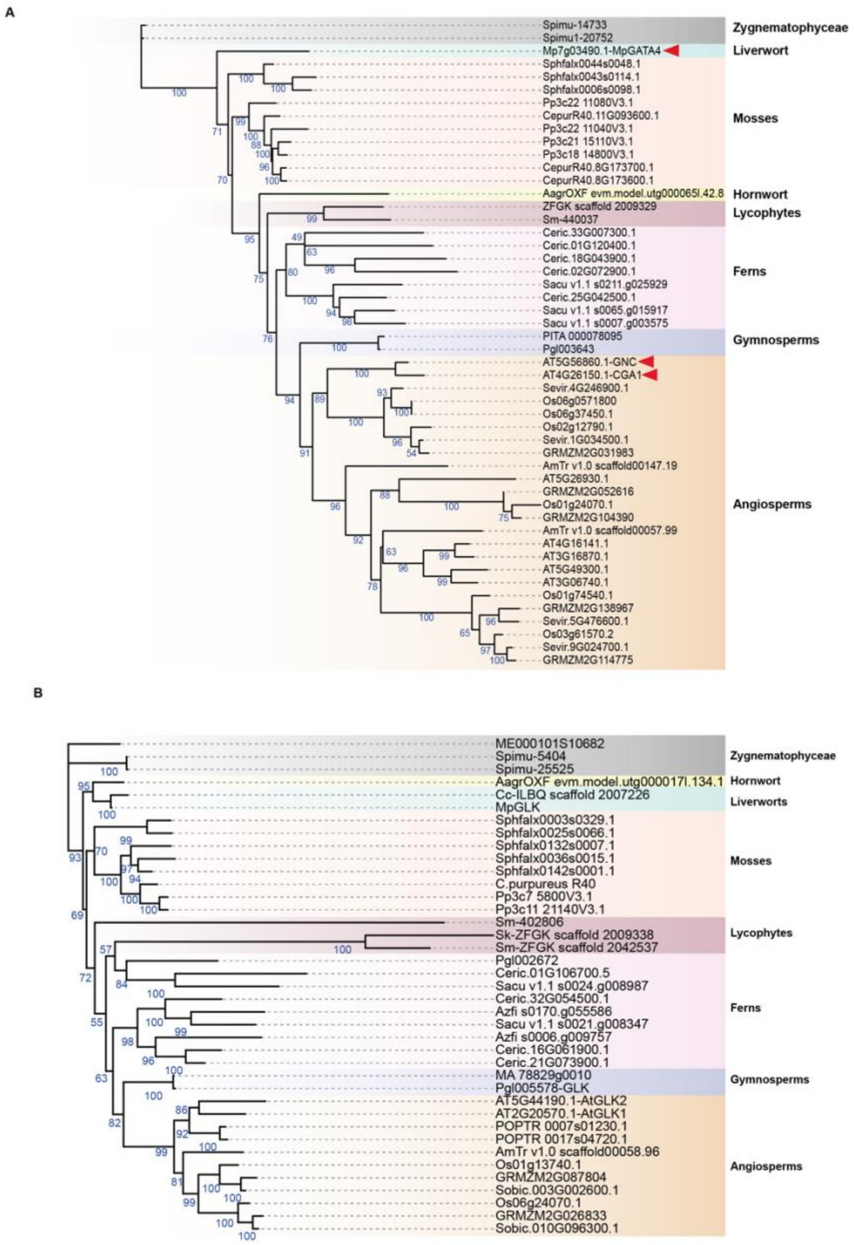


**Figure S3. Phylogenetic analysis of B-Class GATA and GARP-G2-like transcription factors.**

**A)** B-Class GATA phylogeny. Numbers on branches represent ultrafast bootstrap support (Hoang et al. 2018). **B)** GLK phylogeny. Numbers on branches represent ultrafast bootstrap support (Hoang et al. 2018).

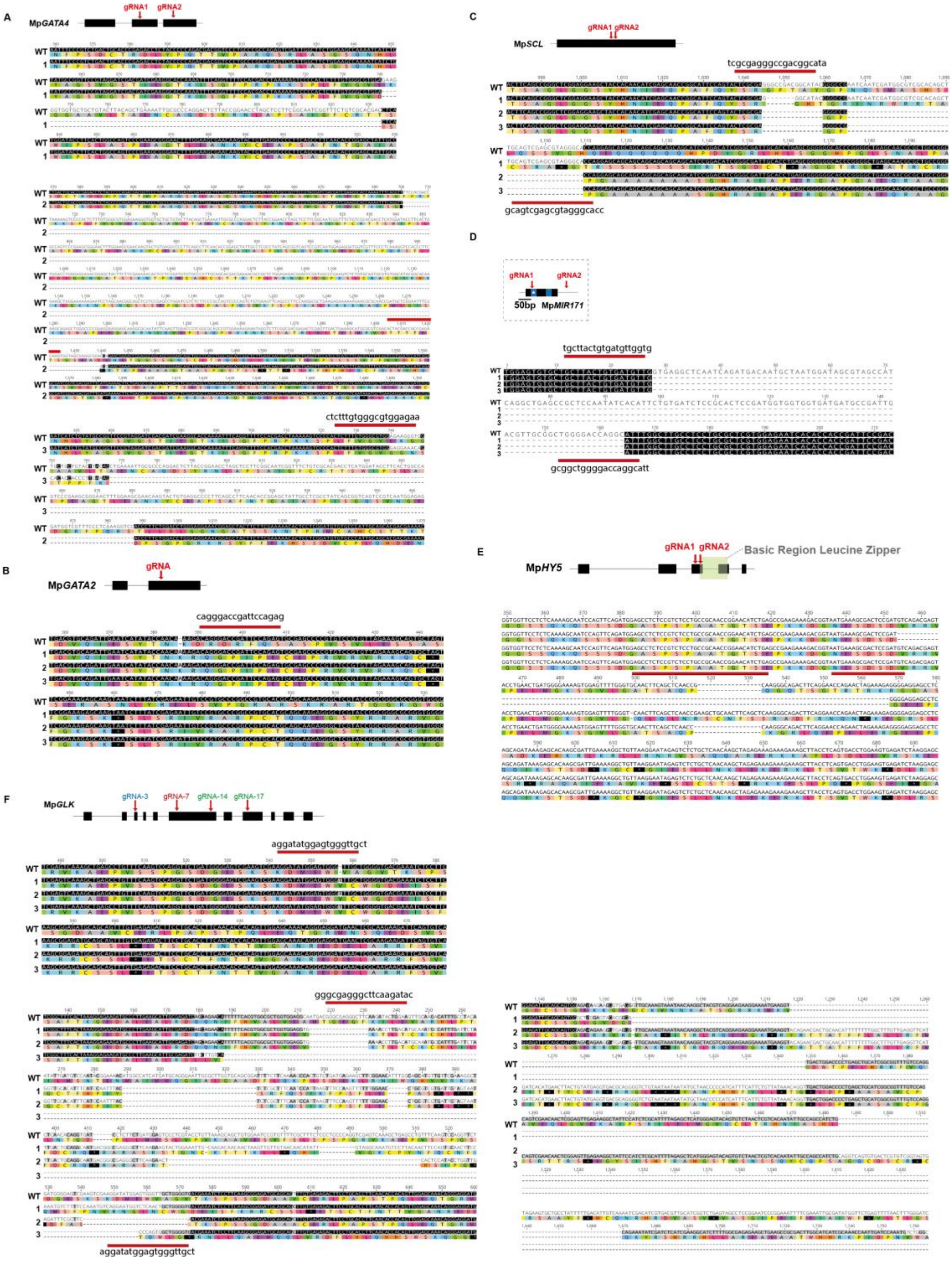


**Figure S4. Sequence analysis of knockout mutant lines.**

**A-E)** For the schematic representation of the Mp*GATA2,* Mp*GATA4,* Mp*SCL,* Mp*MIR171,* Mp*HY5* and Mp*GLK* gene structure exons are shown as black boxes. Position of gRNA(s) is shown with arrow(s). The wild type *M. polymorpha* Cam-1 sequence is shown at the top of the sequence alignment, with the 20 bp gRNA target sequences highlighted with a red line. The amino acid sequence is depicted below the nucleotide sequence.

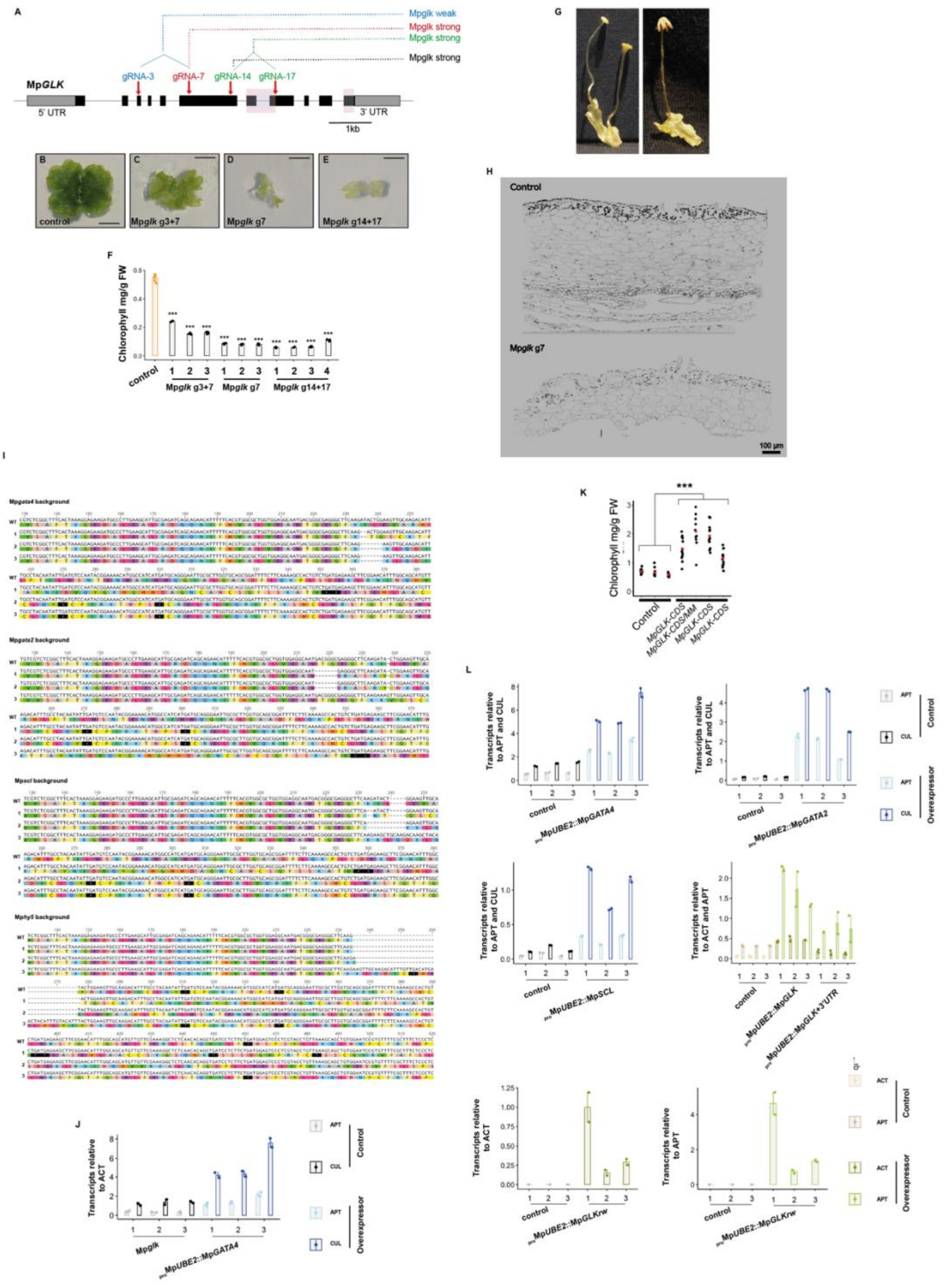


**Figure S5. Analysis of Mp*glk* mutants and quantitative reverse transcription polymerase chain reactions.**

**A)** Schematic representation of Mp*GLK* gene structure showing exons as black rectangles, untranslated regions (UTRs) as grey rectangles and introns as black lines. Characteristic gene domains are highlighted by shaded boxes. Positions of gRNAs used for CRISPR/Cas9 gene editing are shown as red arrows. **B-E)** Images of control and Mp*glk* mutant plants. Scale bars represent 2mm. **F)** Barplots representing chlorophyll content in Mp*glk* mutants. Individual values are shown as dots. Error bars, represent standard deviation of the mean from $n$ = 5. Asterisks indicate statistically significant differences using a two-tailed $t$-test $P \leq 0.0001$ (***),

0.0001 ≤ *P* ≤ 0.001 (**) and n.s.: non-significant. **G)** Representative images of Mp*glk* reproductive organs. **H)** Scanning electron micrograph maps of control (top) and Mp*glk* (bottom) thallus cross sections. **I)** Sequence analysis of Mp*glk,gata4,* Mp*glk,gata2,* Mp*glk,scl* and Mp*glk,hy5* double mutant lines. The wild-type *M. polymorpha* Cam-1 sequence is shown at the top, with the 20 bp gRNA target sequence highlighted with a red line. The amino acid sequence is depicted below the nucleotide sequence.The wild-type *M. polymorpha* Cam-1 sequence is shown at the top. The amino acid sequence is depicted below the nucleotide sequence. **J)** quantitative reverse transcription polymerase chain reaction (qRT-PCR) analysis for Mp*GATA4* over-expression in Mp*glk* mutant plants. *ADENINE PHOSPHORIBOSYL TRANSFERASE 3* (*APT*) and *CULLIN 1* (*CUL*) were used as housekeeping gene controls (Saint-Marcoux et al., 2015). **K)** Chlorophyll content in Mp*GLK* and Mp*GLK/MM* (membrane marker) lines. Jitter plot of chlorophyll content in 'empty vector' controls, Mp*GLK* and Mp*GLK* with membrane markers (Mp*GLK*/MM). Black dots indicate individual transformants, red dots indicate mean. Asterisks indicate statistically significant difference using a two-tailed *t*-test *P* ≤ 0.0001 (***). **L)** qRT-PCR analysis for transgene over-expression confirmation**.** For the qRT-PCR analysis of Mp*GATA4,* Mp*GATA2* and Mp*SCL* over-expression plants, *APT* and *CUL* were used as housekeeping gene controls (Saint-Marcoux et al., 2015). For the qRT-PCR analysis of Mp*GLK* levels in Mp*GLK,* Mp*GLK-CDS+3'UTR* overexpressing lines and Mp*GLK*-CDSrw levels in Mp*GLK*-CDSrw overexpressing lines, *ACTIN 7* (ACT) and *APT* were used as housekeeping gene controls (Saint-Marcoux et al. 2015).

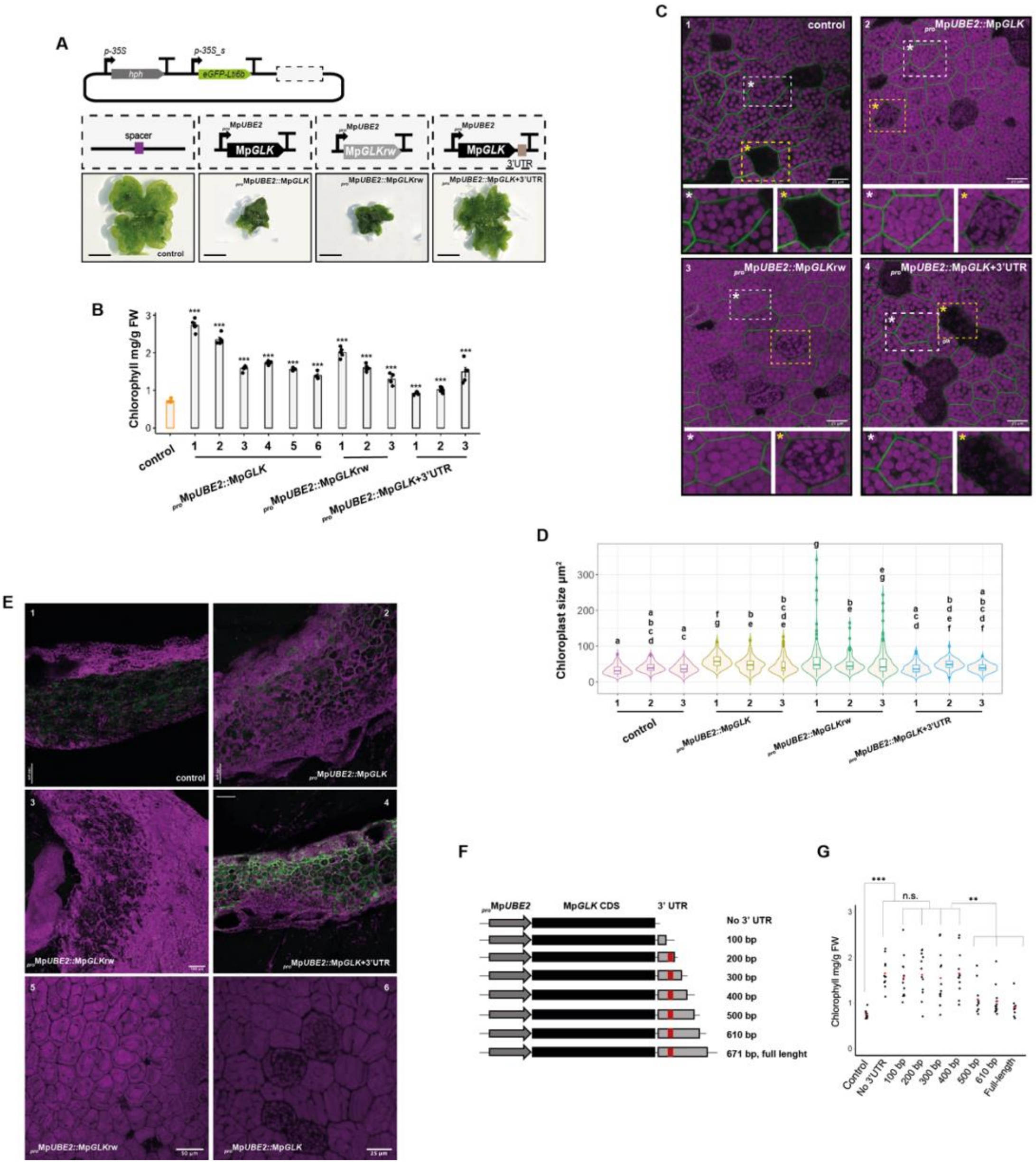

**Figure S6. Mp*GLK* overexpression increases chlorophyll content and Mp*GLK* 3´UTR controls chlorophyll content.**

**A)** Schematic representation of transformation constructs used for Mp*GLK*, Mp*GLK-CDSrw* and Mp*GLK-CDS+3´UTR* overexpression as well as the 'empty vector' control. hph: hygromycin B phosphotransferase, 35S: cauliflower mosaic virus (CaMV) 35S promoter. Bottom: Images of 'empty vector' control plants and plants transformed with Mp*GLK,* Mp*GLK-CDSrw* and Mp*GLK-CDS+3'UTR* overexpression constructs. Scale bars represent 2mm. **B)** Barplots of chlorophyll content in Mp*GLK*, Mp*GLK-CDSrw* and Mp*GLK-CDS+3'UTR* overexpressing lines compared to 'empty vector' controls. Individual values are shown with dots. Error bars, represent standard deviation of the mean from *n* = 3. Asterisks indicate statistically significant difference using a two-tailed *t*-test *P* ≤ 0.0001 (***) and n.s.: non-significant. **C)** Representative confocal microscopy images of control plant gemmae and

Mp*GLK*, Mp*GLK-CDSrw* and Mp*GLK-CDS+3´UTR* overexpressing lines. Chlorophyll autofluorescence is shown in magenta and plasma membrane marked with eGFP shown in green. Magnified cells (white asterisks) and rhizoids precursors (yellow asterisks) are shown under each panel. Scale bars, represent 25 µm. **D)** Boxplot showing chloroplast area range in control versus Mp*GLK*, Mp*GLK-CDSrw* and Mp*GLK-CDS+3´UTR* overexpressing lines. Box and whiskers represent the 25 to 75 percentile and minimum-maximum distributions of the data. Letters show statistical ranking using a *post hoc* Tukey test (with different letters indicating statistically significant differences at P<0.01). Values indicated by the same letter are not statistically different. **E)** Representative confocal microscopy images of thallus sections for control and overexpression of Mp*GLK*, Mp*GLK*-CDSrw and Mp*GLK-CDS+3'UTR* lines. Scale bars, represent 100µm. **F)** Schematic representation of overexpression constructs where Mp*GLK* CDS is fused to a full-length or a series of truncated Mp*GLK* 3´UTR versions. Grey arrows represent Mp*UBE2* promoter, black rectangles - Mp*GLK* CDS, grey rectangles – a full-length or truncated Mp*GLK* 3´UTR. Red boxes indicate Mp*GLK* 3´UTR region harbouring a predicted miR11666.4 and another putative miRNA/siRNA recognition/cleavage sites. **G)** Jitter plot showing chlorophyll content in transgenic lines overexpressing Mp*GLK* CDS with a full-length or truncated Mp*GLK* 3´ UTRs. Black dots represent individual transformants, red dots indicate mean average. Asterisks indicate statistically significant difference using a two-tailed *t*-test $P \leq 0.0001$ (***), $P \leq 0.001$ (**) and n.s.: non-significant.

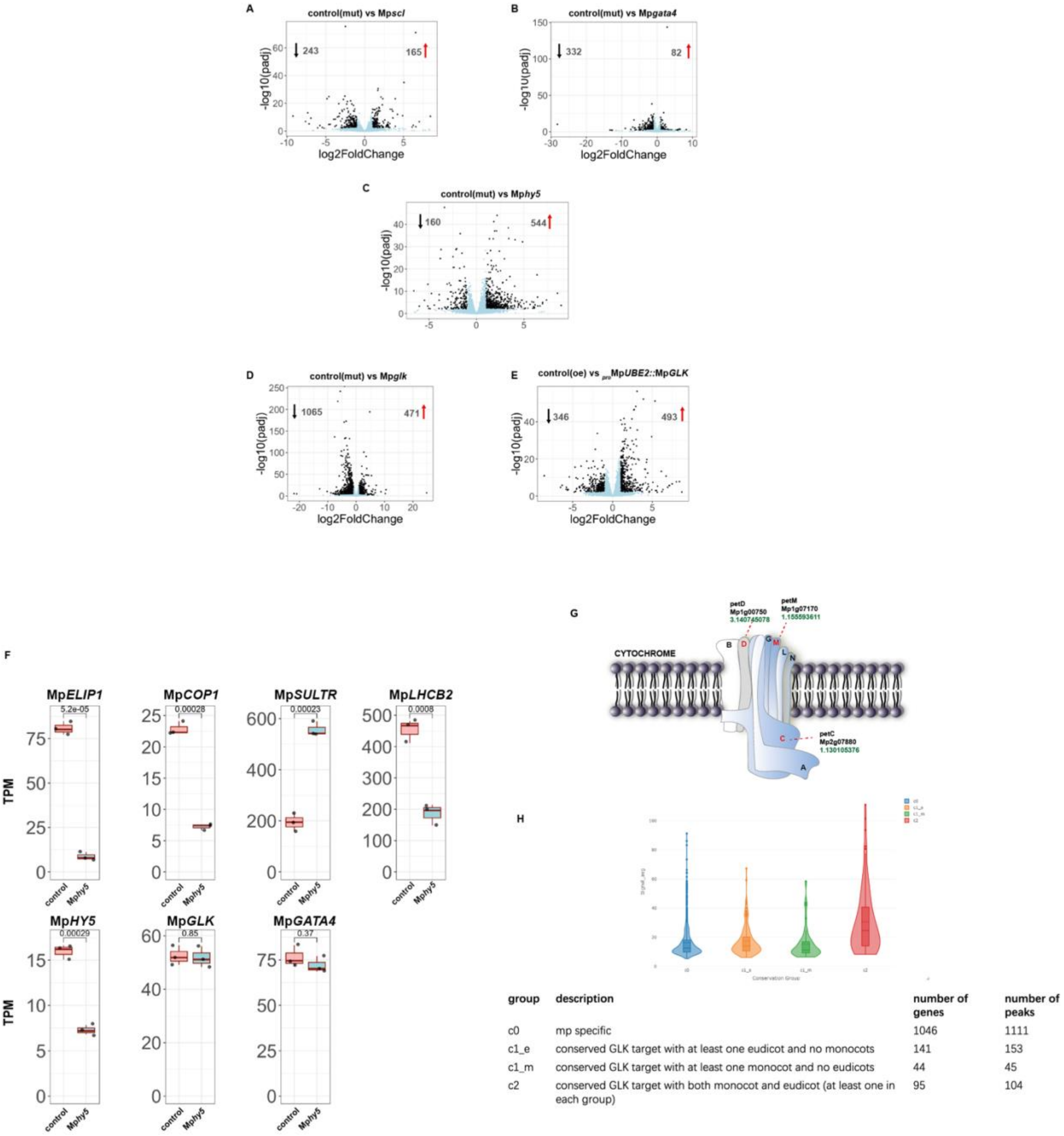


**Figure S7. Expression pattern of M*pGLK* in gemmae and RNAseq gene expression analysis in Mp*scl*, Mp*gata4*, Mp*hy5*, Mp*glk* mutants and Mp*GLK* OE lines.**

**A-E)** Volcano plots showing differentially expressed genes in Mp*scl*, Mp*gata4*, Mp*hy5* and Mp*glk* mutant vs control plants as well as in Mp*GLK* overexpression plants vs control. Blue dots indicate genes with a padj-value > 0.01, while the black ones have a padj-value= < 0.01. The total number of DEGs are indicated at the top of the graph. **F)** Boxplots for representative downregulated genes in Mp*hy5* mutants*. ELIP: Early Light Inducible Protein 1; COP1: Constitutive Photomorphogrnic 1; SULTR: Sulphate Transporter; LHCB2: Light Harvesting Complex B 2.* Data are presented as Transcripts per Million (TPM) values. P-values of two-tailed *t*-test are shown. **G)** Schematic representation of *cytb6f*. Subunits showing an increase (>1) in corresponding transcript levels are highlighted with letters in red. Log2Fold change is shown below the corresponding gene ID. **H)** MpGLK target genes encoding chloroplast-located proteins do not exhibit conservation across these plant species.

**ImageJ macro:**

```
run("Duplicate...");
run("Smooth");
run("Auto Local Threshold", "method=Phansalkar radius=5 parameter_1=0 parameter_2=0 white");
run("Watershed");
run("Analyze Particles...", "size=1-500 circularity=0.01-1.00 display exclude clear add");
```
